# Metabolic and Epigenomic Regulation of Th17/Treg Balance by the Polyamine Pathway

**DOI:** 10.1101/2020.01.23.911966

**Authors:** Chao Wang, Allon Wagner, Johannes Fessler, Julian Avila-Pacheco, Jim Karminski, Pratiksha Thakore, Sarah Zaghouani, Kerry Pierce, Lloyd Bod, Alexandra Schnell, David DeTomaso, Noga Ron-Harel, Marcia Haigis, Daniel Puleston, Erika Pearce, Manoocher Soleimani, Ray Sobel, Clary Clish, Aviv Regev, Nir Yosef, Vijay K. Kuchroo

**Affiliations:** Evergrande center for immunologic diseases, Harvard Medical School and Brigham and Women’s Hospital, Boston, MA 02115, USA; Broad Institute of MIT and Harvard, Cambridge, MA 02142, USA; Department of electrical engineering and computer science, University of California, Berkeley, CA 94720, USA; Center for Computational Biology, University of California, Berkeley, CA 94720, USA; Department of Cell Biology, Harvard Medical School, Boston, MA 02115; Max Planck Institute of Immunobiology and Epigenetics, 79108 Freiburg, Germany; University of Cincinnati, OH, 45221, USA; Palo Alto Veteran’s Administration Health Care System and Department of Pathology, Stanford University School of Medicine, Stanford, CA; Howard Hughes Medical Institute, Department of Biology, Massachusetts Institute of Technology, Cambridge, MA 02140, USA; Koch Institute for Integrative Cancer Research, Massachusetts Institute of Technology, Cambridge, MA 02139, USA; Chan-Zuckerberg Biohub, San Francisco, CA 94158, USA; Ragon Institute of MGH, MIT, and Harvard, Cambridge, MA, USA

## Abstract

Cellular metabolism can orchestrate immune cell function. We previously demonstrated that lipid biosynthesis represents one such gatekeeper to Th17 cell functional state. Utilizing Compass, a transcriptome-based algorithm for prediction of metabolic flux, we constructed a comprehensive metabolic circuitry for Th17 cell function and identified the polyamine pathway as a candidate metabolic node, the flux of which regulates the inflammatory function of T cells. Testing this prediction, we found that expression and activities of enzymes of the polyamine pathway were enhanced in pathogenic Th17 cells and suppressed in regulatory T cells. Perturbation of the polyamine pathway in Th17 cells suppressed canonical Th17 cell cytokines and promoted the expression of Foxp3, accompanied by dramatic shift in transcriptome and epigenome, transitioning Th17 cells into a Treg-like state. Genetic and chemical perturbation of the polyamine pathway resulted in attenuation of tissue inflammation in an autoimmune disease model of central nervous system, with changes in T cell effector phenotype.

## INTRODUCTION

Th17 cells and FoxP3+ regulatory T cells play a key role in maintaining the balance between inflammatory and regulatory functions in the immune system. One key aspect is the balance between Th17 and Treg cells. FoxP3^+^ Tregs play a critical role in maintaining immune tolerance, highlighted by loss-of-function mutations in the Foxp3 gene in human, the master regulator of Tregs, results in the development of IPEX syndrome where patients develop a series of autoimmune pathologies (autoimmune enteropathy, type 1 diabetes, dermatitis) and die prematurely. In contrast, Th17 cells have been shown to be critical for the induction of a number of autoimmune diseases including psoriasis, psoriatic arthritis, ankylosing spondylitis, multiple sclerosis and inflammatory bowel disease [1, 2]. While TGFβ alone can induce FoxP3^+^ Tregs *in vitro*, the addition of proinflammatory cytokine IL-6 suppresses the generation of FoxP3^+^ T cells and together with TGFβ induces generation of Th17 cells. This led to the hypothesis that proinflammatory Th17 and regulatory FoxP3^+^ Tregs are reciprocally regulated, further supported by experiments on the role of these two cytokines in the induction and differentiation of Th17 cells *in vivo* [3–6].

However, not all Th17 cells are pathogenic or disease inducing, and they also play a protective role in mucosal tissues, promoting tissue homeostasis, maintaining barrier function as well as preventing invasion of microbiota at the mucosal sites [7–12]. Th17 cells that are induced by TGFb + IL-6 *in vitro*, produce IL-17 but are not capable of inducing potent tissue inflammation/autoimmunity upon adoptive transfer [13–15]. Additional stimuli, such as IL-1b and IL-23, are needed to evoke pathogenic potential in these Th17 cells [13, 14, 16–20]. Therefore, there appear to be at least two different types of Th17 cells: Th17 cells that are present at homeostasis and do not promote tissue inflammation that we have termed nonpathogenic Th17 cells and the Th17 cells which produce IL-17 together with IFN-g and GMCSF induce tissue inflammation and autoimmunity [21]. Different types of Th17 cells have also been identified in humans where Th17 cells akin to mouse pathogenic Th17 cells have been shown to be specific for *Candida albicans* and non-pathogenic Th17 cells have been shown to be similar to Th17 cells that have specificity for *Staphlococcus aureus* infection [22]. Thus, Treg, non-pathogenic Th17 cells and pathogenic Th17 cells represent a functional spectrum in tissue homeostasis, disease and infection and can be differentiated reciprocally with different cytokine cocktails *in vitro*. However, in addition to cytokines, how these cells are generated *in vivo* and what are the factors that trigger their development of different functional states has not been fully elucidated.

Cellular metabolism is a mediator and modulator of immune cell differentiation and function, which we hypothesized may play a key role in this balance. In a previous study using scRNA-seq of Th17 cells, we identified CD5L as a major regulator that co-varies in its expression with the pro-inflammatory gene module in Th17 cells. Loss of CD5L made Th17 cells highly pathogenic by altering lipid biosynthesis and transcriptional activity of RoR γt, the master transcription factor critical for development and differentiation of Th17 cells [23]. This observation provided a proof of concept that metabolic processes can be directly involved in gene regulation and balancing proinflammatory and regulatory states of Th17 cells.

However, a full appreciation of metabolic circuitry and its connection with immune cell function has been limited by available technologies that typically define the average metabolic state of a large population of cells. We have developed a flux balance analysis algorithm called Compass that allows prediction of metabolic state of a cell using transcriptome data at the single cell level, allowing comprehensive profiling of metabolic pathways even in a smaller number of cells that could not be otherwise interrogated by traditional metabolomic techniques (accompanying manuscript, Wagner *et al*, BioRxiv preprint). Here, we used the Compass algorithm to interrogate the metabolic status of pathogenic and nonpathogenic Th17 cells using scRNA-seq datasets of Th17 cells. We show that enzymes of the polyamine pathway are suppressed and cellular polyamine content is significantly lower in regulatory T cells and non-pathogenic Th17 cells (Th17n) as compared to pathogenic Th17 cells (Th17p) due to alternative fluxing. Perturbation of the polyamine pathway in Th17 cells suppressed canonical Th17 cytokines and promoted Foxp3 expression, shifting the Th17 cell transcriptome in favor of a Treg-like state. We demonstrated that the polyamine pathway is critical in maintaining the Th17-specific chromatin landscape against the induction of Tregs-like program. Consistent with the cellular phenotype, chemical inhibition and genetic perturbation of the polyamine pathway in T cells restricted the development of autoimmune responses in the EAE model.

## RESULTS

### Identifying the polyamine pathway as a candidate in regulating Th17 cell function

To better analyze the metabolic landscape of Th17 cells that may regulate their functional state, we first used two approaches: untargeted metabolomics (Supplemental Figure 1) and standard analysis of single-cell RNAseq data (Figure 1A, B). For both analyses, we compared Th17 cells differentiated from naïve CD4^+^ T cells using two combinations of cytokines: IL-1b+IL-6+IL-23 (Th17p, pathogenic) and TGFb+IL-6 (Th17n, non-pathogenic) that we previously reported to either promote or restrict Th17 cell pathogenicity respectively in the context of the EAE model, and therefore represents the two extremes of functional state of Th17 cells [14, 23]. Untargeted metabolomics identified 1,101 (out of 7,436) metabolic features to be differentially expressed between Th17n and Th17p (BH-adjusted Welch t-test p < 0.05; Supplemental Figure 1). We identified 52 of the differentially expressed metabolites, a third of which (19 / 52) are of lipid nature, consistent with our previous finding that lipid biosynthesis is a key regulator of Th17 cell functions [23], and the rest related to multiple amino-acid pathways. Next, we evaluated the expression of metabolic enzyme genes (“metabolic transcriptome”) of sorted IL-17-GFP+ Th17 cells differentiated *in vitro*, which we previously profiled by scRNA-seq [24]. The distributions of the computational pathogenicity signature scores (computed by expression of key cytokines and transcription factors [14, 24]) of cells from the two conditions were readily distinguishable (p < 3*10^-16^, two-sided Welch t-test). However, there was considerable variation across the individual cells within each condition [23, 24], such that some of the cells from the Th17n condition has higher pathogenicity scores than cells from the Th17p condition (Figure 1A), with a minor distinctive mode of more pathogenic-like cells. This intra-population heterogeneity highlights the benefit of studying the Th17n and Th17p populations at a single-cell level. Interestingly, many of the genes that most covaried with the genes associated with Th17 cell function belonged to ancillary metabolic pathways (Figure 1B), as were most of the identified differentially expressed metabolites, rather than the central and well-studied glycolysis pathway.

**Figure 1.**
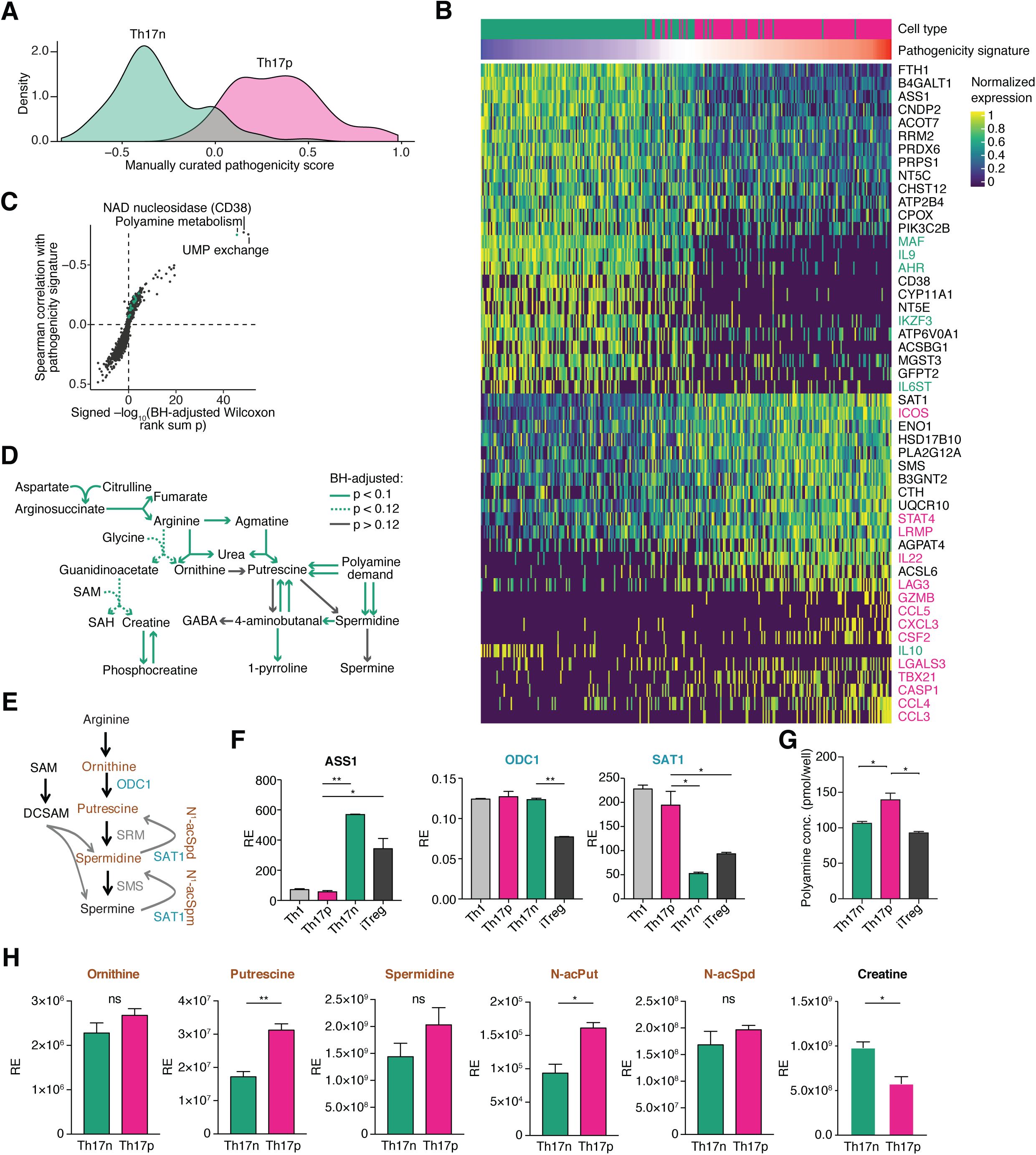
Prediction and metabolic validation of the polyamine pathway as a candidate in regulating Th17 cell function. (**A-B**) Standard single-cell RNAseq analysis of Th17 cells we published in [24]. Briefly, IL-17.GFP+ cells were isolated from pathogenic Th17 cells (Th17p, IL-1b+IL-6+IL-23) and non-pathogenic Th17 cells (Th17n, TGFb+IL-6) differentiated in vitro. **A**, Histogram of a transcriptional pathogenicity score per cell, based on [14]; **B,** Gene expression heatmap of top metabolic genes associated with Th17 cell pathogenicity as we qualified in [14]. Marker genes associated with pro-inflammatory or pro-regulatory programs and used to compute the pathogenicity score are colored in red and green, respectively, black genes are metabolic. Cell are ordered by the ranked pathogenicity score; (**B-C**) Compass analysis of scRNAseq of Th17 cells. **C**, meta-reactions (data-driven clusters reactions with similar Compass scores across cells, with median of two reactions per meta-reaction; Methods) differential activity between Th17p and Th17n conditions is assessed via Benjamini-Hochberg (BH)-adjusted Wilcoxon rank sum p value, signed by the direction of change and by the Spearman correlation of their Compass scores with cell pathogenicity scores, across all cells. Green dots represent meta-reactions containing at least one metabolic reaction that appear in network of panel D. The meta-reaction labelled “polyamine metabolism” contains uptake of putrescine, spermidine, spermidine monoaldehyde, spermine monoaldehyde, and 4-aminobutanal from the extracellular compartment, and the conversion of 4-aminobutanal to putrescine. **D**, a metabolic network that is preferentially active in the pro-regulatory (Th17n) state based on Compass results. Green arrows represent reactions that were predicted to be significantly associated with the Th17n program (BH-adjusted p < 0.1 for their meta-reaction, dashed line for borderline significance, BH-adjusted p < 0.12), grey arrows represent reactions that were not significantly different between Th17p and Th17n; (**E**) Schematic of the polyamine pathway based on KEGG; SAM: S-Adenosyl-Methionine; SAH: S-Adenosyl-Homocysteine. GABA: gamma-aminobutyric acid. (**F-H**) validation of the polyamine pathway. Th17n and Th17p cells are differentiated as in (A) and harvested at 48h for qPCR (F) and 68h for metabolomics (g-h). **F**, qPCR validation of rate-limiting enzymes in polyamine metabolism *ASS1*, *ODC1* and *SAT1*; **G**, Total polyamine content measured by ELISA; **H**, Abundance of metabolites in the polyamine pathway are reported as measured by LC/MS metabolomics.

Next, to obtain a comprehensive view of the metabolic state of each cell despite the inability to measure single cell metabolomic profiles, we investigated the metabolic circuitry of Th17 cells using Compass (accompanying manuscript, Wagner *et al*, BioRxiv preprint; see **Methods**), with the scRNA-seq profiles from sorted IL-17-GFP^+^ Th17 cells [24]. Briefly, Compass is a Flux Balance Analysis (FBA)-based algorithm [25, 26] and utilizes a comprehensive compendium of thousands of metabolic reactions, their stoichiometry, and the enzymes catalyzing them [27]. Compass models *in silico* the fluxes through the network of metabolic reactions, while accounting for the observed expression levels of enzyme-coding transcripts in each cell. It does so by optimizing a series of objective functions, each corresponding to an individual metabolic reaction (rather than a single FBA objective such as biomass production). The result of the optimization procedure is a score for each reaction in each cell, indicative of the potential of the cell to direct flux through that reaction, given the transcriptome of that cell. The Compass scores matrix is then subject to downstream analysis, while relying on the statistical power afforded by scRNA-Seq to derive biological insight from the high-dimensional matrix.

Analysis of the Compass scores for each reaction across all single cells in our data (Figure 1C) showed that among those metabolic reactions significantly correlated with Th17 cell pathogenicity, the polyamine pathway stood out as one that is differentially activated in pathogenic *vs*. nonpathogenic Th17 cells (Figure 1C and **Supplemental Table 1**). To explore this, we constructed a data-driven metabolic network anchored around putrescine, the entrance metabolite into the canonical polyamine synthesis, by including adjacent metabolites whose reactions are predicted to be negatively associated with pathogenicity (Figure 1D). While several polyamine-associated genes (*e.g.*, *Sat1* in Figure 1B) are differentially expressed between Th17p and Th17n, the network tied the differential polyamine metabolism to differences in upstream and downstream metabolic reactions which could not be captured from differential gene expression directly. Specifically, Compass predicted that Th17n cells are more active in arginine metabolic pathways, lying upstream of putrescine, and in alternative fates of putrescine (other than conversion to spermidine, along the canonical polyamine synthesis pathway) (Figure 1D). We hypothesized that the arginine/polyamine pathway may be a metabolic bifurcation point that can regulate Th17 cell function and set out to investigate this metabolic network surrounding polyamines.

### Cellular polyamines are suppressed in regulatory T cells and nonpathogneic Th17

To investigate the polyamine metabolic process (Figure 1E), we first asked whether critical enzymes of this pathway are differentially expressed in different CD4^+^ T cell subsets using qPCR. Ornithine Decarboxylase 1 (ODC1) and Spermidine/Spermine N1 Acetyltransferase 1 (SAT1) are the rate-limiting enzymes of polyamine biosynthesis and catabolic processes, respectively, and Ornithine decarboxylase antizyme 1 (OAZ1) can regulate the enzymatic activity of ODC1. ODC1 catalyzes ornithine to putrescine, the first step of the polyamines biosynthesis; SAT1 regulates the intracellular recycling of polyamines and their transport out of the cell. *SAT1*, but not *ODC1* or OAZ1, was suppressed in Th17n *vs*. Th17p cells (Figure 1F and data not shown). Intriguingly, both ODC1 and SAT1 expression was lower in Tregs, whereas Ass1, an enzyme upstream of the polyamine biosynthesis pathway is upregulated, consistent with Compass-predicted alternative flux in the polyamine neighborhood (Figure 1F). Collectively, these data suggest the polyamine pathway may be associated with functional state beyond Th17 cells.

As the polyamine pathway, similar to most metabolic pathways, is regulated beyond the transcriptional level, we next directly measured total cellular polyamine content using an enzymatic assay (**Methods**). Compared to Th17p cells, T_regs_ and Th17n cells have significantly reduced levels of total polyamines (Figure 1G), reflective of either reduced import, biosynthesis or increased export of polyamines in these cells.

To further investigate the concentrations and activities of different polyamines in Th17 cells at different functional states, we applied both targeted metabolomics and carbon tracing. We differentiated Th17n and Th17p cells for 68 hours (**Methods**) and measured the amount of polyamines and related precursors in cell and media by LC/MS (Figure 1H and S1B). Consistent with Compass’s predictions, there was higher creatine content in Th17n *vs*. Th17p cells. On the other hand, while the total amount of cellular ornithine, precursor to polyamines, was comparable between Th17n and Th17p cells, there was a significant increase of putrescine and acetyl-putrescine content in Th17p cells (Figure 1H), indicative of increased activity of this pathway in Th17p cells, consistent with the enzymatic assay. Of note, cellular spermidine (or acetyl-spermidine) content was not different between the conditions, and spermine was not detected (Figure 1H). The reduced putrescine and its acetyl form in Th17n cells are not due to increased export, as we observed very little polyamines in the media in either Th17n or Th17p cells (Supplemental Figure 1B). These data suggest that polyamines accumulate within Th17p cells and the main function of SAT1 in Th17p cells may be to recycle rather than to export polyamines.

To directly investigate polyamine biosynthesis, we cultured differentiated Th17n and Th17p cells in the presence of low amount of carbon or hydrogen labeled arginine or citrulline, which can be used to synthesize ornithine, precursor to the polyamine pathway (Supplemental Figure 1C, D). First, we harvested cells and media for LC/MS at 0, 1, 5 and 24 hours post addition of arginine. While there was comparable accumulation of labeled cellular guanidinoacetic acid, a byproduct of arginine conversion into ornithine, in Th17n and Th17p cells over time (Supplemental Figure 1C), Th17p cells accumulated higher intracellular amounts of putrescine, acetylputrescine and acetylspermidine, consistent with increased polyamine biosynthesis and/or recycling activity in these cells (Supplemental Figure 1C). Conversely, there were higher levels of labeled arginine in Th17n cells vs. Th17p cells, prompting us to investigate whether Th17n cells can better synthesize (as opposed to better uptake) arginine, which would be consistent with increased ASS1 expression (Figure 1F) in these cells. To this end, we harvested cells for LC/MS 24 hours after addition of labeled citrulline, a precursor to arginine synthesis. Indeed, there was higher accumulation of labeled arginine in Th17n cells (Supplemental Figure 1D). Collectively, our targeted metabolomics and carbon tracing data suggest that Th17n cells accumulate arginine, consistent with Compass’s prediction (Figure 1D), and that Th17p cells preferentially synthesize or recycle polyamines. We conclude that differences in the alternative flux hinged on polyamine biosynthesis is associated with the different functional states of Th17 cells.

### ODC1 or SAT1 inhibition restricts Th17 cell function in a putrescine-dependent manner

To investigate the functional relevance of these metabolic changes, we studied the effects of polyamine pathway inhibitors on differentiation of pathogenic and nonpathogenic Th17 cells *in vitro*, using previously defined culture conditions. We first used difluoromethylornithine (DFMO), an irreversible inhibitor of ODC1 (Figure 2A), the enzyme that catalyzes the conversion of ornithine to putrescine. Enzymatic assays of *in vitro* differentiated Th17n, Th17p, or iTreg cells treated by DFMO comfirmed its suppression of polyamines in all three cell types (Supplemental Figure 2A). At an optimized concentration where we observed similar viability between control and treatment, DFMO significantly inhibited IL-17 expression in both Th17n and Th17p cells by intracellular staining and flow cytometry (Figure 2B), as well other canonical Th17 cytokines such as IL-17A, IL-17F, IL-21 and IL-22, while promoting IL-9 expression in supernatant from both Th17n and Th17p cultures (Figure 2C). DFMO did not consistently influence, IFNg, TNFa, IL-13, IL-10 or IL-5 expression (Supplemental Figure 2B). IL-17 inhibition does not appear to be solely related to regulation of IL-2 production [28], as DFMO promoted IL-2 expression in supernatant from only Th17p, but not Th17n cells (Figure 2C). Polyamines can influence cell proliferation. While we did observe reduced cell proliferation in cultures treated with DFMO, the frequency of IL-17^+^ cells was significantly reduced in cells that have divided just once (data not shown), suggesting DFMO can regulate Th17 cell function independent of cellular proliferation. The increase in IL-9 following DFMO treatment also supports the hypothesis that DFMO is not universally inhibiting viability of Th17 cells and enhances Th9 derived cytokines.

**Figure 2.**
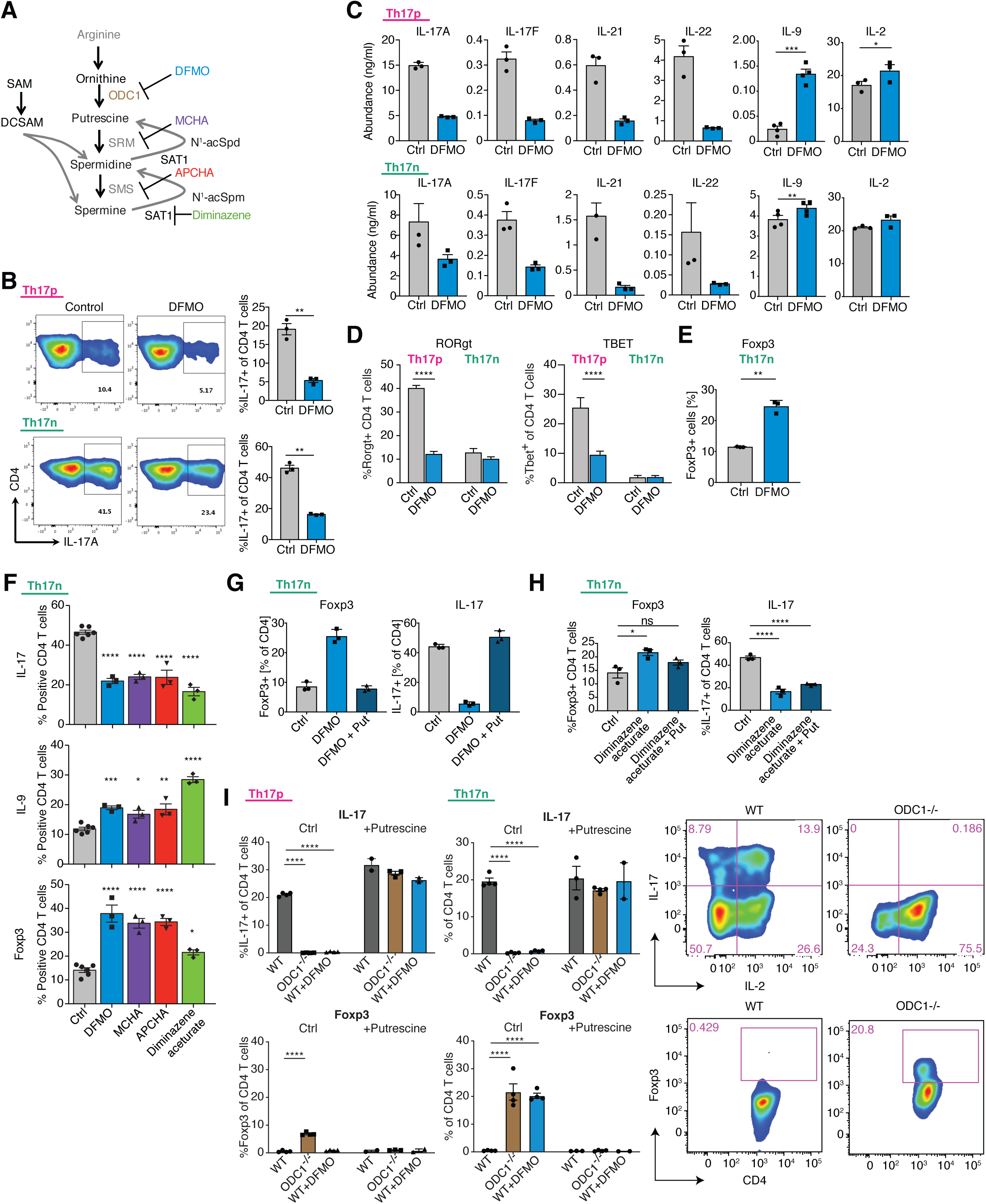
Chemical and genetic interference with the polyamine pathway suppress canonical Th17 cell cytokines. **(A)**, Polyamine pathway overview depicting inhibitors of ODC1 (DFMO), SRM (MCHA), SMS (APCHA) and SAT1 (Diminazene aceturate). (**B-E**) The effects of DFMO on Th17n and Th17p cells differentiated as in Figure 1. DFMO were added at the time of differentiation cytokines. All analysis performed on day 3. **B-C,** Flow cytometric analysis of intracellular cytokines (B) and secreted cytokines by legendplex (C); **D**, Flow cytometric analysis of transcription factor expression in Th17n and Th17p; **E**, Flow cytometric analysis of Foxp3 expression in Th17n. (**F**) Comparison of IL-17A, IL-9 and FoxP3 expression following treatment with Ctrl, DFMO, MCHA, APCHA or Diminazene aceturate in *in vitro* differentiated Th17n cells. (**G-H**) The rescue effect of adding putrescine on inhibitors to ODC1 (G) or SAT1 (H). Expression of IL-17A and FoxP3 following combinational treatment of DFMO (G) or Diminazene aceturate (H) with or without the addition of 2.5mM Putrescine. (**I**) The effects of genetic perturbation of ODC1. Th17n and Th17p cells are generated from naïve T cells isolated from WT or ODC1^-/-^ mice and treated with control or DFMO in combination with 0 or 2.5mM Putrescine. Flow cytometric analysis of intracellular IL-17 and Foxp3 are shown. Each dot represents biological replicates performed with different mice. All statistical analyses are performed using pair-wise comparison or one-way anova.

To determine whether DFMO inhibited Th17 cell differentiation, we measured the expression and activity of key transcription factors. Interestingly, DFMO suppressed Rorgt and Tbet expression in Th17p but not Th17n cells (Figure 2D), suggesting a nuanced effect. Consistently, DFMO decreased pStat3, and not total Stat3 protein levels, only in Th17p but not Th17n cells (Supplemental Figure 2C). IL-17 inhibition is not due to increased Foxo1 activity, another critical regulator of Th17 cell function, as DFMO promoted pFoxo1(S256) in both types of Th17 cells, which would have resulted in a net increase in IL-17 expression (Supplemental Figure 2C). Given the known reciprocal relationship between Th17 cells and Tregs, and as DMFO also impacted polyamine concentration in Tregs, we asked whether DFMO can regulate Foxp3 expression in Th17 cells, even under Th17 differentiation conditions. We observed increased frequency of Foxp3^+^ cells in Th17n but not Th17p conditions (Figure 2E), presumably because TGFb is required for the differentiation in this condition and DFMO strengthens TGFb derived activity to induce T_reg_ differentiation over Th17 cells.

To determine whether other enzymes of the polyamine pathway could play a similar role in regulating Th17 cell function, we used inhibitors of spermine synthase (SRM), spermidine synthase (SMS), and SAT1 (Figure 2A). Similar to DFMO, inhibitors of any of the polyamine biosynthesis enzymes resulted in suppression of IL-17 and upregulation of IL-9 and Foxp3 expression, the latter in Th17n cells (Figure 2F). Furthermore, inhibiting SAT1 by diminazene, a rate-limiting enzyme of polyamine acetylation and recycling, had similar effects to DFMO (Figure 2F). SAT1 perturbation was previously reported to have a feedback effect on ODC1 activity and vice versa [29–31]. Consistent with this finding, inhibition with DFMO consistently suppressed SAT1 expression in both Th17n and Th17p cells (Supplemental Figure 2D). Thus, it may be the flux of polyamines and not metabolites per se that modulate Th17 cell function.

Finally, we confirmed that the effect of DFMO is through the inhibition of ODC1, as addition of putrescine to cells treated with DFMO completely reversed their phenotype (Figure 2G). Interestingly, addition of putrescine during SAT1 inhibitor treatment also partially reversed the upregulation of Foxp3, but not suppression of IL-17 (Figure 2H), suggesting putrescine flux may be particularly important in the control of the regulatory program in Th17 cells. Overall, the inhibitor data are consistent with a role of the polyamine pathway in regulating Th17 cell differentiation, but genome-wide profiling would be necessarily to further support this claim.

### ODC1^-/-^ Th17 cells promoted Foxp3 expression

To further confirm the effects of chemical inhibition of polyamine pathway on Th17/Treg differentiation, we tested the impact of genetic perturbation of ODC1 on the differentiation and functions of Th17 cells, using cells isolated from WT and ODC1^-/-^ mice. Similar to DFMO treatment, there was complete inhibition of Th17 canonical cytokines, such as IL-17A, IL-17F and IL-22, but not IFNg, in ODC1^-/-^ Th17 cells (Figure 2I upper panel and S2E). ODC1 deficiency did not lead to a decrease in Rorgt expression (data not shown), but there was a dramatic loss of Th17 canonical cytokines, consistent with loss of the Th17 program. Furthermore, ODC1^-/-^ Th17n cells upregulated Foxp3 expression, consistent with promotion of a T_reg_ program (Figure 2I, lower panel). Finally, all the observed effects of ODC1^-/-^ were rescued by addition of putrescine (Figure 2I and S2E).

### DFMO restricts Th17-cell transcriptome and epigenome in favor of Treg-like state

To gain mechanistic insight on the effects of inhibiting polyamine biosynthesis in Th17 cells, we profiled by RNA-Seq Th17n, Th17p, and iTreg cells treated with DFMO or control. DFMO had a profound impact on the transcriptome of all Th cell lineages, driving Th17 cells towards Treg cell profiles in Principal Components Analysis (PCA) (Figure 3A, PC1). To gain further insights, we determined the aggregate effect of DFMO on genes up-regulated (n= 1,284), down-regulated (n= 1,255) or comparable (n= 8,257) in Th17 *vs*. T_reg_ cells (Figure 3B). In both Th17n and Th17p cells, DFMO suppressed the Th17 cell specific gene set, and promoted the Treg-specific transcriptome (Figure 3C, **Supplemental Table 2 and 3**). Specifically, canonical Th17 cell genes such as *Il17a, Il17f* and *Il23r* were significantly suppressed, whereas Treg related genes, such as Foxp3, were upregulated (Figure 3B). There was no significant effect of DFMO treatment on genes expressed comparably in Th17 cells and Treg, nor did DFMO have an effect in Treg cells (Figure 3B **and** 3C). These results are consistent with a model where the polyamine pathway is important for restricting the iTreg-like program in Th17 cells at both functional states (Figure 3A).

**Figure 3.**
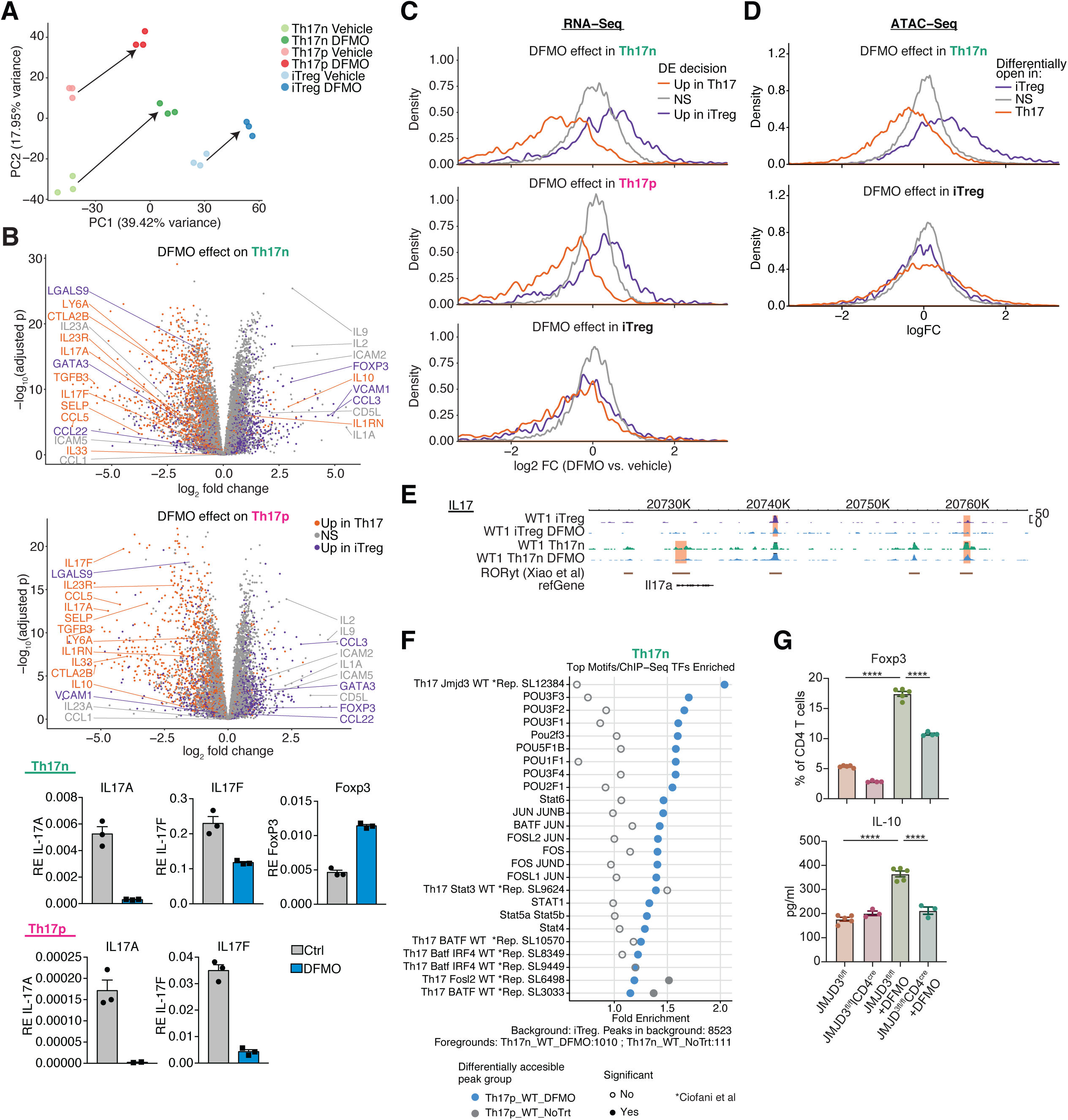
DFMO treatment promotes Treg-like transcriptome and epigenome. (**A-C**) Th17n, Th17p and iTreg cells were differentiated and harvested at 68h for live cell sorting and population RNAseq. **A**, PCA plot showing in vitro differentiated Th17n (green), Th17p (red) and iTregs (blue) in the presence (lighter color) or absence of DFMO. **B**, Volcano plots (upper) and qPCR validation (lower) showing affected genes by DFMO treatment in Th17n and Th17p cells. Th17 (orange) and iTreg specific genes (violet) are highlighted. **C,** Histograms showing the effects of DFMO on iTreg, Th17n and Th17p transcriptome. Transcriptome space is divided into those up-regulated in Th17 cells (orange), Treg (violet) or neither (grey). (**D-F**) Th17n and iTreg cells were differentiated and harvested at 68h for live cell sorting and population ATAC-seq. **D**, Histograms showing the effects of DFMO on chromatin accessibility as measured by ATAC-seq. The accessibility regions are divided into those more accessible in Th17 cells (orange), Treg (violet) or neither (grey). **E**, IGV plots of *Il17* regions. Regions significantly altered (DESeq2, BH-adjusted p <0.05) by DFMO treatment and binding sites for RORγt [32] are highlighted. **F**, Motif enrichment analysis of *in vitro* differentiated Th17n in the presence (red) or absence (green) of DFMO for iTreg specific genes. (**G**), Cells were cultured under Th17n condition as in Figure 2 with DFMO or solvent control (water), replated to rest at 68h in new plate and harvested at 120h for analysis of intracellular Foxp3 expression and IL-10 expression in supernatant.

The profound impact of DFMO on the transcriptome prompted us to investigate the mechanism by which the polyamine pathway regulates Th17 cell functions. As DFMO does not appear to consistently restrict phosphorylation of key Th17 cell regulators, particularly not in Th17n cells (Supplemental Figure 2C), we hypothesized that polyamines may impact the epigenome. Consistent with a role of the polyamine pathway in affecting chromatin modification, we observed significant changes in expression of many chromatin modifiers (Supplemental Figure 3A).

To test this hypothesis, we measured chromatin accessibility by ATAC-seq in Th17n and iTregs cells treated with either control or DFMO (**Methods**). Overall, DFMO treatment resulted in considerable changes in accessible peaks in both types of Th cells (Supplemental Figure 3B **and Supplemental Table 4A and 4B**). Next, we asked whether DFMO preferentially altered accessibility to regions specific to Th17 cells and iTregs. To this end, we partitioned all accessible peaks into (1) those more accessible in Th17 cells (n = 10,431), (2) more accessible in iTregs (n = 3,421), and (3) comparably accessible in both (n = 34,591) (Figure 3D, **Supplemental Table 3, and Methods**). Consistent with the expression changes, following DFMO treatment there was a significant shift towards less accessibility in Th17 specific regions and more accessibility in Treg specific regions (Figure 3D and **Supplemental Table 3**). Differentially accessible regions were found near loci encoding key effector molecules (**Supplemental Table 4A and 4B**). For instance, DFMO treatment significantly restricted peaks in the promoter and intergenic regions of *Il17a-Il17f* that corresponds to Rorgt binding site (using ChIP-seq data from [32]) known to regulate IL17 expression (Figure 3E). Thus, DFMO treatment can shape chromatin accessibility in favor of an iTreg epigenomic landscape, and this may contribute to the emergence of iTreg transcriptional program in DFMO-treated Th17 cells.

### The chromatin regulator JMJD3 partially mediates the polyamine-dependent effect on Th17 cells

To investigate which transcription factors (TFs) may be responsible for the suppression of the Th17 specific program and upregulation of the iTreg program, we looked for putative binding sites (based on DNA binding motifs or ChIP-seq data when available) that significantly overlap with regions whose accessibility is modulated by DFMO (Figure 3F and **Supplemental Table 5**). We restricted our analysis to genomic regions that are typically accessible only in Tregs (compared to Th17 cells) and may be modulated by DFMO (Figure 3F and Supplemental Figure 3C). In Th17n cells, DFMO increased accessibility near potential binding sites of the chromatin regulator JMJD3 along with a number of POU-domain containing TFs.

As JMJD3 is a known regulator of T cell plasticity [33], we tested whether it also contributes to the genome-wide shifts induced by DFMO. We analyzed the effect of DFMO on Th17 cells differentiated from naïve CD4 T cells isolated from control or JMJD3^fl/fl^CD4^cre^ mice (Figure 3G). Supporting our hypothesis, the upregulation of Foxp3 by DFMO in Th17n cells was partially abrogated in the absence of JMJD3, and loss of JMJD3 also reduced the DFMO-dependent upregulation of IL-10 in Th17n cells (Figure 3G).

### Perturbation of ODC1 and SAT1, key enzymes of the polyamine pathway, alleviates EAE

To investigate the relevance of the polyamine pathway *in vivo*, we took two approaches to perturbing it in the context of CNS autoimmune disease, EAE: chemical inhibition of ODC1 and T-cell specific genetic deletion of SAT1 (Figure 4). As targeting multiple nodes in the polyamine pathway resulted in upregulation of Foxp3 during Th17 differentiation *in vitro* (Figure 2 and **3**), we hypothesized that targeting rate-limiting enzymes in polyamine pathway *in vivo* would regulate induction of EAE.

**Figure 4.**
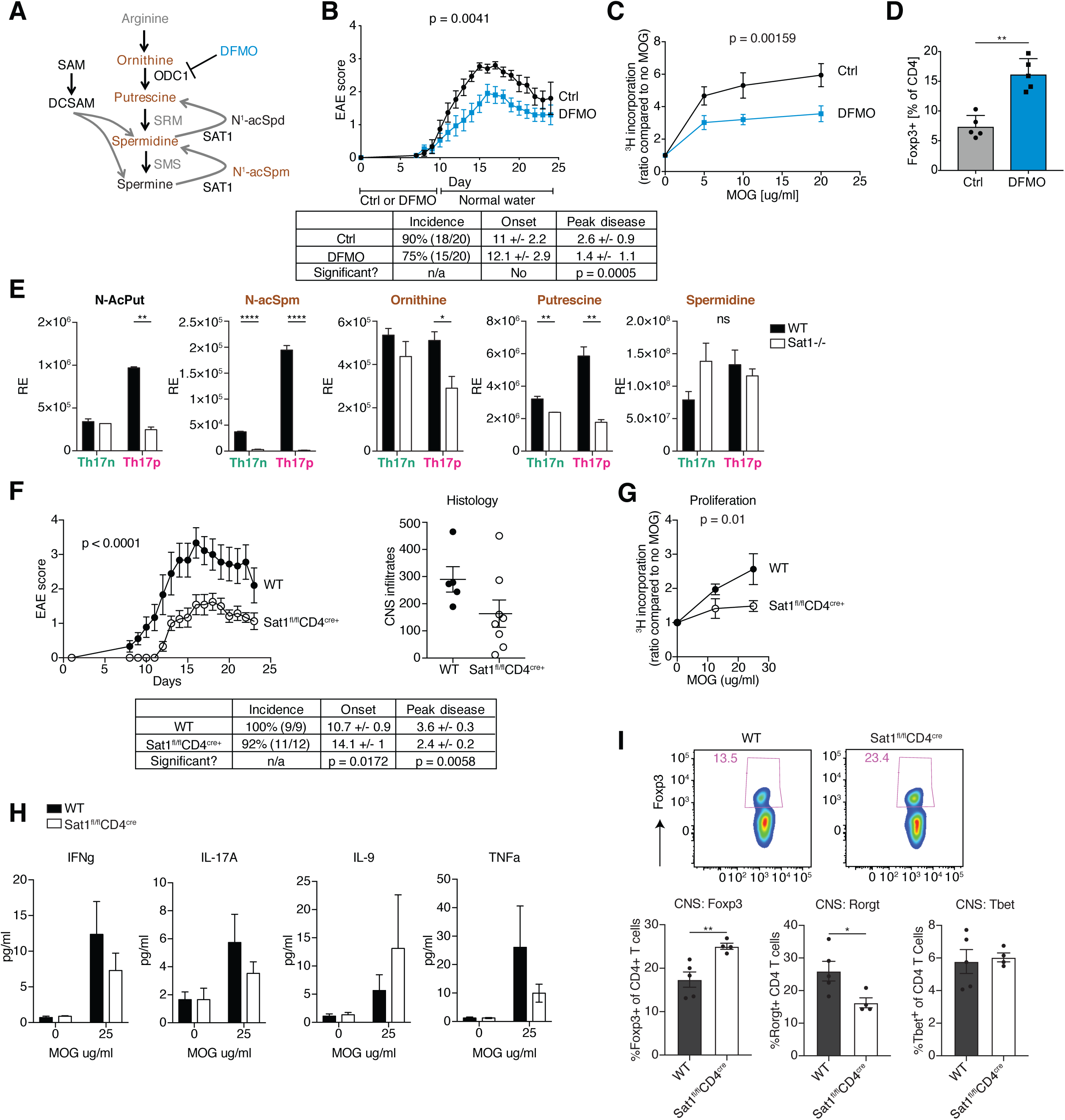
Targeting ODC1 and SAT1 alleviate EAE. (**A**) Schematics of the polyamine pathway; (**B-D**) The effects of chemical inhibition of ODC1 by DFMO on EAE. Wildtype mice were immunized with MOG_35-55_/CFA to induce experimental autoimmune encephalomyelitis and followed for clinical scoring. DFMO were provided in drinking water from day 0 for 10 days in experimental group. **B**, Clinical score over time. Graph shows pooled results from 3 independent experiments. **C**, Antigen-specific cell proliferation is measured by thymidine incorporation after culturing cells isolated from draining lymph node of mice (d15 post immunization) with increasing dose of MOG_35-55_ peptide for 3 days. **D**, Flow cytometric analysis of intracellular Foxp3 expression in T cells isolated from CNS at day 15 post immunization. (**E-I**) The effects of genetic perturbation of SAT1. **E**, The effects of SAT1 deficiency on metabolome. Abundance of metabolites in the polyamine pathway were determined by LC/MS based metabolomics. Th17n and Th17p cells were differentiated in vitro from naïve cells isolated from WT or SAT1^-/-^ mice. (**F-I**) The effects of SAT1 deficiency on EAE. EAE were induced as in (B) in wildtype and SAT1^fl/fl^CD4^cre^ mice. **F**, Clinical score (left) and histological score (right) showing the number of CNS infiltrates show representative experiment with 5 WT and 8 CKO mice. Similar results were obtained from 2 independent experiments summarized in table (lower). **G-H**, Cells were isolated from draining lymph node of mice (d23 post immunization) and co-cultured with increasing dose of MOG_35-55_ peptide for 3 days. Antigen-specific cell proliferation is measured by thymidine incorporation (**G**) and antigen-specific cytokine secretion by legendplex (**H**). **I**, Flow cytometry analysis of intracellular transcription factor expression in CD4 T cells isolated from CNS day 15 post immunization. Linear regression analysis (b, c, f, g), two-way anova (e) and student *t* test (d, i) were used for statistical analysis.

We first tested ODC1 inhibition by adding DFMO in the drinking water for mice immunized with MOG/CFA for the induction of EAE (**Methods**). DFMO significantly delayed the onset and severity of EAE (Figure 4B). Consistently, we observed a significantly reduced antigen-specific recall response in the draining lymph node of DFMO treated animals (Figure 4C). Further analysis of lymphocytes isolated from the CNS showed no difference in the frequency of cytokine producing cells, but increased frequency of Foxp3^+^ CD4^+^ T cells out of all CD4 T cells (Figure 4D and data not shown), consistent with the polyamine biosynthesis pathway as an important positive regulator of the balance between proinflammatory Th17 cells and Foxp3^+^ Tregs and induction of autoimmune CNS inflammation, which is highly dependent on Th17 cells.

Since administering DFMO in the drinking water could affect multiple cell types, we also genetically deleted SAT1, the rate limiting enzyme of the polyamine pathway, in CD4^+^ T cells (SAT1^fl/fl^CD4^cre^). We confirmed that genetic deletion of SAT1 in T cells resulted in loss of polyamine acetylation as reflected in reduced levels of acetyl-putrescine and acetyl-spermidine (Figure 4E). Notably, loss of SAT1 also resulted in reduced level of putrescine in Th17 cells, likely through a feedback mechanism. This is consistent with reports in other cell types [31] and our *in vitro* inhibitor data (Supplemental Figure 2), suggesting similar effect of DFMO and SAT1 deletion in the context of T cell biology may be due to overall changes in polyamine flux. Indeed, we observed significantly delayed onset and severity of EAE in SAT1^fl/fl^CD4^cre^ mice (Figure 4F). Similar to global inhibition of ODC1 by DFMO treatment, we observed an inhibition of antigen-specific recall responses as measured by T cell proliferation (Figure 4G). Although we did not observe significant differences in cytokine production (Figure 4H and Supplemental Figure 4A), there was a trend towards a decrease in IFN-g, IL-17 and TNF production with an increase in IL-9 production in response to antigen (Figure 4H). Furthermore, there was a significant increase in the proportion of Foxp3^+^CD4^+^ T cells (out of all CD4 T cells) and concomitant decrease of Rorgt^+^CD4^+^ T cells isolated from the target organ (CNS) of SAT1^fl/fl^CD4^cre^ mice (Figure 4I). Notably, the frequencies of Foxp3^+^ or Rorgt^+^ cells are not different in the draining lymph node (Supplemental Figure 4B), suggesting that the effect of SAT1 on T cells may be amplified in tissue recall responses. Thus, using both chemical and genetic perturbations at multiple levels, we demonstrated that the polyamine pathway is an important mediator of autoimmune inflammation.

## DISCUSSION

To understand the functional relevance of metabolic pathways in Th17 cells, we utilized metabolomics, a novel computational algorithm (Compass, companion manuscript) and chemical and genetic perturbation to investigate the functional metabolic networks that impact Th17 pathogenicity. In this study, we investigated in depth the metabolic circuitry centered around the polyamine pathway. We demonstrated that 1) At the transcriptome level, Compass points to the significance of the polyamine pathway as a top candidate in association with Th17 cell pathogenicity and implicates reactions upstream and downstream of putrescine to be associated with functional phenotype of Th17 cells; 2) As predicted by Compass and measured by enzymatic assay and LC/MS metabolomics, we showed that Th17 cells at different functional state have alternative metabolic flux anchored around arginine and putrescine, the precursor to polyamines, and that both regulatory T cells and non-pathogenic Th17 cell have reduced cellular content of polyamines; 3) Chemical targeting of multiple enzymes in the polyamine pathway and genetic deletion of ODC1 resulted in suppression of the Th17 functional program and upregulation of Foxp3 in a putrescine dependent manner; 4) Inhibiting polyamine biosynthesis shifts Th17 cells in favor of Treg-like transcriptome and epigenome; 5) Targeting ODC1 and SAT1 both resulted in upregulation of Foxp3 *in vivo* and inhibition of effector Th17 cells and regulation of EAE. Taken together, we have provided evidence supporting a critical role of the polyamine pathway in suppressing regulatory program in Th17 cells.

Th17 cells are critical in inducing autoimmune inflammation. In fact, loss of all the components in Th17 pathway including TGF-b, IL-6, IL-1 or IL-23 results in inhibition of Th17 differentiation, upregulation of FoxP3+ Tregs and suppression of EAE. Because of reciprocal generation of Tregs vs. Th17 cells, the effects observed with the inhibition of polyamine pathway may be unique to the diseases where Th17 cells are the effector cells. Whether the effect of polyamine pathway can be generalized to other autoimmune diseases (e.g. autoimmune colitis or type 1 diabetes), where Th1 or NK cells are the effectors, need to be further evaluated. In fact the effects of blocking polyamine pathway in diverting Th17 differentiation to Treg phenotype was much more profound in generating nonpathogenic Th17 (differentiation with TGFb) than in pathogenic Th17 cells (differentiation with IL-1b and IL-23). This observation suggests that inhibition of the pathway may have an effect that is unique to Th17 driven diseases.

The significance of the polyamine pathway in autoimmune diseases is further supported by anecdotal data that polyamine levels are increased in several autoimmune diseases [34, 35] and it is thought that aberrant polyamine metabolism can contribute to autoantigen stabilization [36]. Here we present a potential mechanism of how the polyamine pathway can regulate Th17/Treg balance and impact development of autoimmunity. DFMO is an FDA-approved drug for cancer therapy. We showed that DFMO has significant impact in curtailing EAE, providing the grounds/mechanism for drug repurposing. It should be noted that while targeting any enzyme in the polyamine pathway resulted in similar effects in Th17 cells in vitro, genetic manipulation of ODC1 and SAT1 are not identical in that while both ODC1 and SAT1 deletion promoted Foxp3 expression (Figure 2I and data not shown), ODC1 but not SAT1 suppressed Th17 cytokine expression *in vitro* (Figure 2I and data not shown). Further studies are necessary to understand the mechanistic difference within the polyamine pathway.

By studying the metabolic differences within the same lineage of effector Th17 cells, we unexpectedly uncovered a central role of the polyamines in regulating Th17-Treg balance. This study suggests a functional role of metabolic pathways beyond energy production. One of the observations made in this study is the role that polyamine pathway plays in shaping the epigenetic landscape of differentiating immune cells. In fact, looking at the ATACseq and RNAseq profiles of Th17 cells activated in the presence of inhibitors of the polyamine pathway shows profound global ATACseq changes concomitantly with changes in transcription, differentiation and function. Polyamines appear to regulate gene expression, cell proliferation and stress responses due to their ability to bind to nucleic acids (both DNA, RNA), alter posttranslational modification and regulate ion channels [37, 38]. A number of studies have suggested the role of polyamines in regulating gene expression due to their polycationic nature and ability to function as a sink to S-adenosylmethionine and Acetyl-coA, both critical metabolites for histone modifications [29, 30, 39, 40]. Furthermore, intracellular polyamines and their analogues are also known to inhibit lysine-specific demethyltransferases such as LSD1 [41] and thereby changing epigenetic landscape affecting development and differentiation. Thus, it stands to reason that metabolic processes that impact polyamines will not only affect energetics but more broadly including shaping the epigenome and transcriptome by the resultant metabolites that are produced during the process of development or differentiation. In this vein, a number of developmental disorders (eg. Snyder-Robinson syndrome) have been associated with maladapted polyamine metabolism [42].

It is very clear that when immune cells take up residence in different tissues they also change their transcriptomes and attain specialized or different functions. Notable examples of this issue has been shown in tissue Tregs [43] and macrophages [44], where the cells look very different transcriptomically depending on the tissue of residence. We and others have observed a similar situation in Th17 cells, where they differ in their function of whether they are in lymph nodes, gut or CNS, as observed by the scRNAseq analysis of Th17 cells [23, 24]. Based on our studies, presented here, we suggest that the metabolic activity of the cell within a defined tissue may have a profound impact in the epigenome and transcriptome, resulting in their changed or specialized functions. With the emerging cell atlases and mapping transcriptome of tissues resident immune cells at the single cell level, the Compass algorithm will provide a powerful tool for studying metabolic pathways across different cell types in different tissues, taking advantage of the wealth of single cell data sets that are being published.

In summary, our study highlights the advantage of utilizing single cell genomics and novel algorithms in studying cellular metabolism, providing roadmaps for studying metabolic pathways in immune cells across normal or diseased tissues. The study validates the predictions made by algorithms, both in vitro and in vivo and shows that interfering with these metabolic pathways identified by Compass have profound effect on the function of the effector cells, by regulating both epigenome and transcriptome of the Th17 cell.

## ACKNOWLEDGEMENTS

This investigation is supported in part by a Career Transitional Fellowship from the National Multiple Sclerosis Society awarded to CW. NY and AW were supported by the Chan Zuckerberg Biohub. JF was supported by a Max Kade fellowship awarded by the Austrian Acadamy of Science (ÖAW). VKK is supported by grants from National Institutes of Health (R01NS045937, RO1 NS 30843, R01AI144166, P01AI073748, P01AI039671 and P01AI056299)

## AUTHOR CONTRIBUTIONS

CW, AW, AR, NY and VKK conceptualized the study. CW conceived and designed experiments with help from JF; CW, AW, JAP, CC conceived the metabolomics experiments; CW, JF and SZ performed most of the experiments; JAP and KP performed LC/MS metabolomics; AW and JK analyzed RNAseq and ATACseq datasets; PT provided help with ATACseq experiments; PT and AS optimized ATAC-seq protocol, with input from AR; NR, MH provided critical assay support; DP, EP, MS provided critical materials; CW, VKK and NY supervised the study. CW, AW, AR, NY and VKK wrote the manuscript with contributions from all authors.

## METHODS

### Mice

C57BL/6 wildtype (WT) were obtained from Jackson laboratory (Bar Harbor, ME). SAT1flox mice were kindly provided by Dr. Manoocher Soleimani (University of Cincinnati), which we crossed to CD4cre to generate conditional T cell deletion of SAT1. Note that only male mice were used in all experiments as SAT1 is an X chromosome gene and female mice have incomplete deletion due to random inactivation of x chromosome. ODC1^fl/fl^CD4^cre^ were gifted by Dr. Erika Pearce (Max Planck Institute). For experiments, mice were matched for sex and age, and most mice were 6–10 weeks old. Littermate WT or Cre-mice were used as controls. All experiments were conducted in accordance with animal protocols approved by the Harvard Medical Area Standing Committee on Animals or BWH IACUC.

### T cell differentiation culture and flow cytometry

Naïve CD4+CD44-CD62L+CD25-T cells were sorted using BD FACSAria sorter and activated with plate-bound anti-CD3 (1µg/ml) and antiCD28 antibodies (1µg/ml) in the presence of cytokines at a concentration of 5 × 10^5^ cells/ml. For T cell differentiations the following combinations of cytokines were used: pathogenic Th17: 25ng/ml rmIL-6, 20ng/ml rmIL-1b (both Miltenyi Biotec) and 20ng/ml rmIL-23 (R&D systems); non-pathogenic Th17: 25ng/ml rmIL-6 and 2ng/ml of rhTGFb1 (Miltenyi Biotec); iTreg: 2ng/ml of rhTGFb1; Th1: 20ng/ml rmIL-12 (R&D systems); Th2: 20ng/ml rmIL-4 (Miltenyi Biotec). Intracellular cytokine staining was performed after incubation for 4-6h with Cell Stimulation cocktail plus Golgi transport inhibitors (Thermo Fisher Scientific) using the BD Cytofix/Cytoperm buffer set (BD Biosciences) per manufacturer’s instructions. Transcription factor staining was performed using the Foxp3/Transcription Factor Staining Buffer Set (eBioscience). Proliferation was assessed by staining with CellTrace Violet (Thermo Fisher Scientific) per manufacturer’s instructions. Apoptosis was assessed using Annexin V staining kit (BioLegend). Phosphorylation of proteins to determine cell signaling was performed with BD Phosflow buffer system (BD bioscience) as per manufacturer’s instructions.

### Inhibitors and metabolites

For differentiation experiments, cells were harvested at 72 hours and were performed in the presence or absence of 100-200µM DFMO, 500µM trans-4-Methylcyclohexylamine (MCHA, both Sigma), 500µM N-(3-Aminopropyl)cyclohexylamine (APCHA, Santa Cruz Biotechnology), 50µM Diminazene aceturate (Dize, Cayman Chemical) with or without 2.5 mM Putrescine (Sigma, P7505) as indicated.

### Compass analysis

Compass is descried in detail in a companion manuscript (Wagner et al). In the following we provide a high level description of the algorithm.

Compass integrates scRNA-Seq profiles with prior knowledge of the metabolic network to infer a metabolic state of the cell. The metabolic network we use here consists of 7,440 reactions and 2,626 metabolites (Recon2 database, [27]), along with reaction stoichiometry, gene-enzyme-reaction associations and biochemical constraints (such as reaction irreversibility and nutrient availability).

Compass builds on the paradigm of Flux Balance Analysis (FBA) to model metabolic fluxes, namely the rate by which chemical reactions convert substrates to products [25, 26, 45, 46] (Orth, Thiele, and Palsson 2010; O’Brien, Monk, and Palsson 2015; Lewis, Nagarajan, and Palsson 2012; Palsson 2015). The modeling is based on linear programming, maximizing a certain objective (here, flux through a given reaction), while using the metabolic network to pose constraints.

In its first step, Compass is agnostic to any measurement of gene expression levels and computes, for every metabolic reaction *r*, the maximal flux 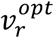 it can carry without imposing any constraints on top of those imposed by stoichiometry and mass balance. Next, Compass assigns every reaction in every cell a penalty inversely proportional to the mRNA expression associated with its enzyme(s) in that cell. Compass then finds a flux distribution which minimizes the overall penalty incurred in any given cell *i* (summing over all reactions), while maintaining a flux of at least 0.95 ⋅ 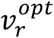 in *r*. The Compass score of reaction *r* in cell *i* is the negative of that minimal penalty (so that lower scores correspond to lower potential metabolic activity). Intuitively, these scores reflect how well adjusted is each cell’s transcriptome to maintaining high flux through each reaction. To reduce the effects of data sparsity (characteristic of scRNA-Seq) Compass uses an information-sharing approach. Instead of treating each cell in isolation, the score vector for each cell is determined by a combined objective that balances the effects in the cell in question with those in its ten nearest neighbors (based on similarity of their RNA profiles).

After applying Compass to the scRNA-Seq of Th17 cells, we aggregated reactions that were highly correlated across the entire dataset (Spearman rho ≥ 0.98) into meta-reactions (with median of two reactions per meta-reaction) for downstream analysis. For our ranking analysis in Figure 1C, we prioritized meta-reactions with differential predicted activity between the Th17p and Th17n conditions. To this end, we used Wilcoxon’s rank sum p-values (comparing Th17 cells differentiated under the non-pathogenic conditions vs. Th17 cells differentiated under the pathogenic conditions) and Spearman rank correlation (correlating reaction scores with pathogenicity scores across cells).

### qPCR

RNA was isolated using RNeasy Plus Mini Kit (Qiagen) and reverse transcribed to cDNA with iScript cDNA Synthesis Kit (Bio-Rad). Gene expression was analyzed by quantitative real-time PCR on a ViiA7 System (Thermo Fisher Scientific) using TaqMan Fast Advanced Master Mix (Thermo Fisher Scientific) with the following primer/probe sets: *Ass1 (*<u>Mm00711256_m1</u>), *Odc1* (Mm02019269_g1), *Sat1* (Mm00485911_g1), *Srm* (Mm00726089_s1), *Sms (*<u>Mm00786246_s1</u>), *Il-17a* (<u>Mm00439618_m1</u>), *Il-17f* (<u>Mm00521423_m1</u>), *Foxp3* (<u>Mm00475162_m1</u>), *Tead1* (Mm00493507_m1), *Taz* (Mm00504978_m1), and *Actb* (Applied Biosystems). Expression values were calculated relative to *Actb* detected in the same sample by duplex qPCR.

### Polyamine ELISA

Cell pellets of *in vitro* differentiated cells were frozen down and further processed with the Total Polyamine Assay Kit (BioVision Inc.) according to the manufacturer’s instructions.

### Metabolomics / Carbon tracing

For untargeted metabolomics, Th17 cells were differentiated as described. Culture media were snap frozen. Cells were harvested at 96h. 1×10^6^ cells per sample were snap frozen and extracted in either 80% methanol (for fatty acids and oxylipids) or isopropanol (for polar and nonpolar lipids). Two liquid chromatography tandem mass spectrometry (LC-MS) methods were used to measure fatty acids and lipids in cell extracts.

For carbon tracing experiments Th17 cells were differentiated as described. At 48h, cells were washed and cultured in media supplemented with Arginine (^13^C6, Sigma, Cat# 643440) or aspartic acid (^13^C4, Sigma, Cat# 604852) for 1, 5 and 24 hours.

### Legendplex

Cytokine concentrations in supernatants of *in vitro* cultures were analyzed by the LegendPlex Mouse Th Cytokine Panel (13-plex) (BioLegend) according to the manufacturer’s instructions and analyzed on a FACS LSR II (BD Biosciences).

### RNA-seq

For population (bulk) RNA-seq, *in vitro* differentiated T-cells were sorted for live cells and lysed with RLT Plus buffer and RNA was extracted using the RNeasy Plus Mini Kit (Qiagen). Full-length RNA-seq libraries were prepared as previously described [47] and paired-end sequenced (75 bp × 2) with a 150 cycle Nextseq 500 high output V2 kit.

### Bioinformatic analysis of RNA-seq data

Alignment, quantification, and computation of pathogenicity signatures based on single-cell transcriptomes were conducted as described in the accompanying manuscript (Wagner *et al*, BioRxiv preprint). Briefly, raw scRNA-seq reads from Gaublomme et al. (2015) [24] (Figure 1) were aligned with Bowtie2, quantified into TPM gene expression with RSEM. Quality control tested and batch effects and other nuisance factors removed with SCONE [48].

To compute a pathogenicity score for each cell we used a similar scheme as in [24]: For each cell we take the average z-scored normalized log expression of pro-pathogenic markers (CASP1, CCL3, CCL4, CCL5, CSF2, CXCL3, GZMB, ICOS, IL22, IL7R, LAG3, LGALS3, LRMP, STAT4, TBX21) and of pro-regulatory markers (AHR, IKZF3, IL10, IL1RN, IL6ST, IL9, MAF), with the latter multiplied by −1.

Bulk RNA libraries from DFMO-or vehicle-treated Th17p, Th17n, or Treg were studied with 3 replicates per condition for a total of 18 libraries as shown in Figure 3A. Genes that are associated with a Th17 or Treg programs (orange and purple, respectively, in Figure 3B-C) were determined by differential expression test between bulk RNA libraries of (vehicle-treated) Th17n and Th17p on one side and Treg on the other with BH-adjusted p ≤ 0.05 and absolute value of log2 fold-change of at least 1.5. Genes associated with the Th17p or Th17n program (magenta and green, respectively, in Supplemental Figure 3A) were determined by differential expression test between bulk RNA libraries of (vehicle-treated) Th17p vs. Th17n with the same thresholds. The PCA shown in Figure 3A was computed on the set of 3,414 that were differentially associated with Th17, Treg, Th17p, or Th17n programs by the aforementioned criteria to focus it on the subspace of the transcriptome relevant to Th17 pathogenicity phenotypes.

### ATAC-seq

For population ATAC-seq, *in vitro* differentiated T-cells were sorted for live cells and stored in Bambanker freezing media (Thermo Fisher Scientific) at −80°C until further processing. Prior to library preparation, cells were thawed at 37 °C and washed with PBS. For ATAC-seq, cell pellets were lysed and tagmented in 1X TD Buffer, 0.2ul TDE1 (Illumina), 0.01% digitonin, and 0.3X PBS in 40ul reaction volume following the protocol described in [49]. Transposition reactions were incubated at 37 °C for 30 min at 300 rpm. The DNA was purified from the reaction using a MinElute PCR purification kit (QIAGEN). The whole resulting product was then PCR-amplified using indexed primers with NEBNext High-Fidelity 2X PCR Master Mix (NEB). First, we performed 5 cycles of pre-amplification. We sampled 10% of the pre-amplification reaction for SYBR Green quantitative PCR to assess the number of additional cycles needed for final amplification. After purifying the final library with the MinElute PCR purification kit (QIAGEN), the library was quantified for sequencing using qPCR and a Qubit dsDNA HS Assay kit (Invitrogen). Libraries were sequenced on an Illumina NextSeq 550 system with paired-end reads of 37 base pairs in length.

### Alignment of ATAC-Seq and Peak Calling

All ATAC-Seq reads were trimmed using Trimmomatic [50] to remove primer and low-quality bases. Reads < 36bp were dropped. Reads were then passed to FastQC [http://www.bioinformatics.babraham.ac.uk/projects/fastqc/] to check the quality of the trimmed reads. The paired-end reads were then aligned to the mm10 reference genome using bowtie2 [51], allowing maximum insert sizes of 2000 bp, with the “--no-mixed” and “--no-discordant” parameters added. Reads with a mapping quality (MAPQ) below 30 were removed. Duplicates were removed with PicardTools, and the reads mapping to the blacklist regions and mitochondrial DNA were also removed. Reads mapping to the positive strand were moved +4 bp, and reads mapping to the negative strand were moved −5bp following the procedure outlined in [52] to account for the binding of the Tn5 transposase.

Peaks were called using macs2 on the aligned fragments [53] with a qvalue cutoff of 0.001 and overlapping peaks among replicates were merged.

### Tests of Differential Accessibility

Differential accessibility was assessed using DESeq2 [54] on with a matrix of peaks (merging all samples) by samples. Similar to common practice in the analysis of differential gene expression, our analysis of differential accessibility was conducted using the number of observed Tn5 cuts (i.e., number of reads).

Peaks that are associated with a Th17 or Treg programs (orange and purple, respectively, in Figure 3D) were determined by differential accessibility test between libraries of (vehicle-treated) Th17n and Th17p on one side (unpublished dataset) and Treg on the other with BH-adjusted p ≤ 0.05 and absolute value of log2 fold-change of at least 1.

### Reprocessing of published ChIP-Seq data

ChIP-Seq Peaks from Xiao et al 2014 [32] were transferred from mm9 to mm10 using the UCSC liftOver tool. ChIP-Seq replicates from Ciofani et al 2012 were downloaded and were trimmed using Trimmomatic [26] to remove primer and low-quality bases. Reads were then passed to FastQC [http://www.bioinformatics.babraham.ac.uk/projects/fastqc/] to check the quality of the trimmed reads. These single-end reads were then aligned to the mm10 reference genome using bowtie2 [27], allowing maximum insert sizes of 2000 bp, with the “--no-mixed” and “--no-discordant” parameters added. Reads with a mapping quality (MAPQ) below 30 were removed. Duplicates were removed with PicardTools, and the reads mapping to the blacklist regions and mitochondrial DNA were also removed.

ChIP-Seq peaks were called in each replicate, versus a control sample, using macs2 [29] with a qvalue cutoff of 0.05.

### Enrichment of motifs and ChIP-seq peaks in differentially accessible regions

Peaks were considered differentially accessible if they had a BH-adjusted p <0.05. We calculated fold enrichment of various genomic features in these peaks (described below) versus a background set of peaks. q-values were estimated using q-value package. [Storey JD, Bass AJ, Dabney A, Robinson D. *qvalue: Q-value estimation for false discovery rate control*. http://github.com/jdstorey/qvalue]

### Motifs / Annotation Tracks

PWM’s for motifs were downloaded from the 2018 release of JASPAR [55, 56]. We used fimo [56] to identify motifs in mm10, and applied the default threshold of 1e-4. We also included regulatory features from the ORegAnno database[57], (iii) conserved regions annotated by the multiz30way algorithm, and repeat regions annotated by RepeatMasker (http://www.repeatmasker.org).

### GREAT Pathways / Genes

Loci were associated with pathways using GREAT[58], submitted with the rGREAT package (https://github.com/jokergoo/rGREAT). We retrieved pathways found in the MSigDB Immunologic Signatures, MSigDB Pathway, and GO Biological Process databases. Loci were mapped to genes using GREAT.

### Experimental Autoimmune Encephalomyelitis (EAE)

For active EAE immunization, MOG35-55 peptide was emulsified in complete freund adjuvant (CFA). Equivalent of 40µg MOG peptide was injected per mouse subcutaneously followed by pertussis toxin injection intravenously on day 0 and day 2 of immunization. Mice were treated with 0.5% DFMO in drinking water for 10 days as indicated. DFMO was replenished every third day.

### Statistical Analysis

Unless otherwise specified, all statistical analyses were performed using the two-tail student t test using GraphPad Prism software. P value less than 0.05 is considered significant (P < 0.05 = *; P < 0.01 = **; P < 0.001 = ***) unless otherwise indicated.

## Supplementary FIGURE LEGENDS

**Supplemental Figure 1.**
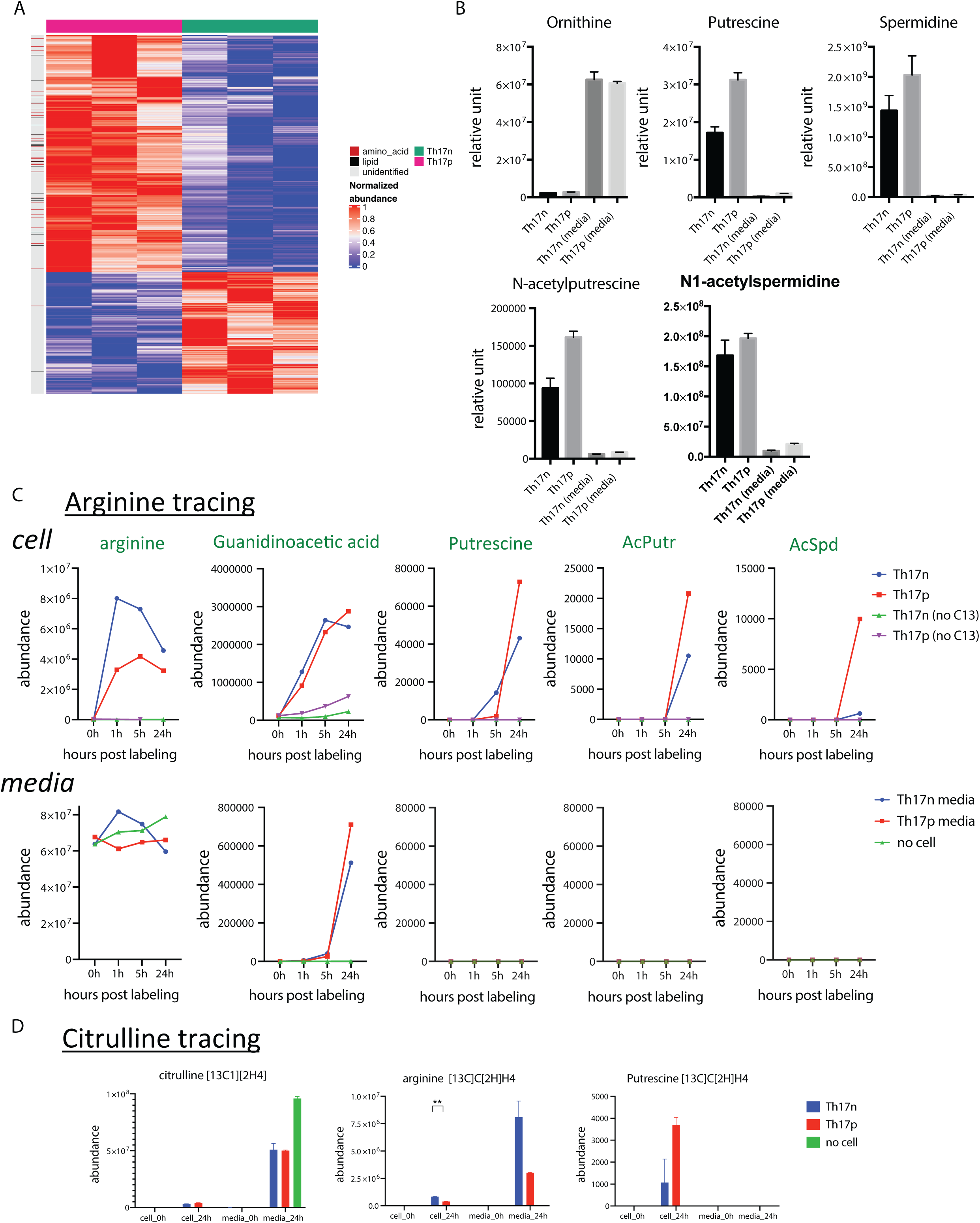
Prediction and metabolic validation of the polyamine pathway as a candidate in regulating Th17 cell function. (**A**) Metabolomics analysis of Th17n (green) and Th17p (red) cells. Cells were differentiated as described (Methods) and harvested at 68h for LC/MS based metabolomics. Shown are 1,101 differentially expressed metabolites between Th17n and Th17p (BH-adjusted Welch t-test p < 0.05), 52 of which are identified and divided between lipids and amino-acid derivatives; (**B**) Metabolomics analysis of the polyamine pathway as in Figure 1H. Cell lysates as well as media from Th17n and Th17p differentiation cultures are shown. (**C-D**) Carbon tracing in the polyamine pathway. Th17n and Th17p cells were differentiated as described (Methods), lifted to rest at 68 hours and pulsed with C13 labeled Arginine (C) or Citrulline (D) followed by LC/MS analysis at time points indicated.

**Supplemental Figure 2.**
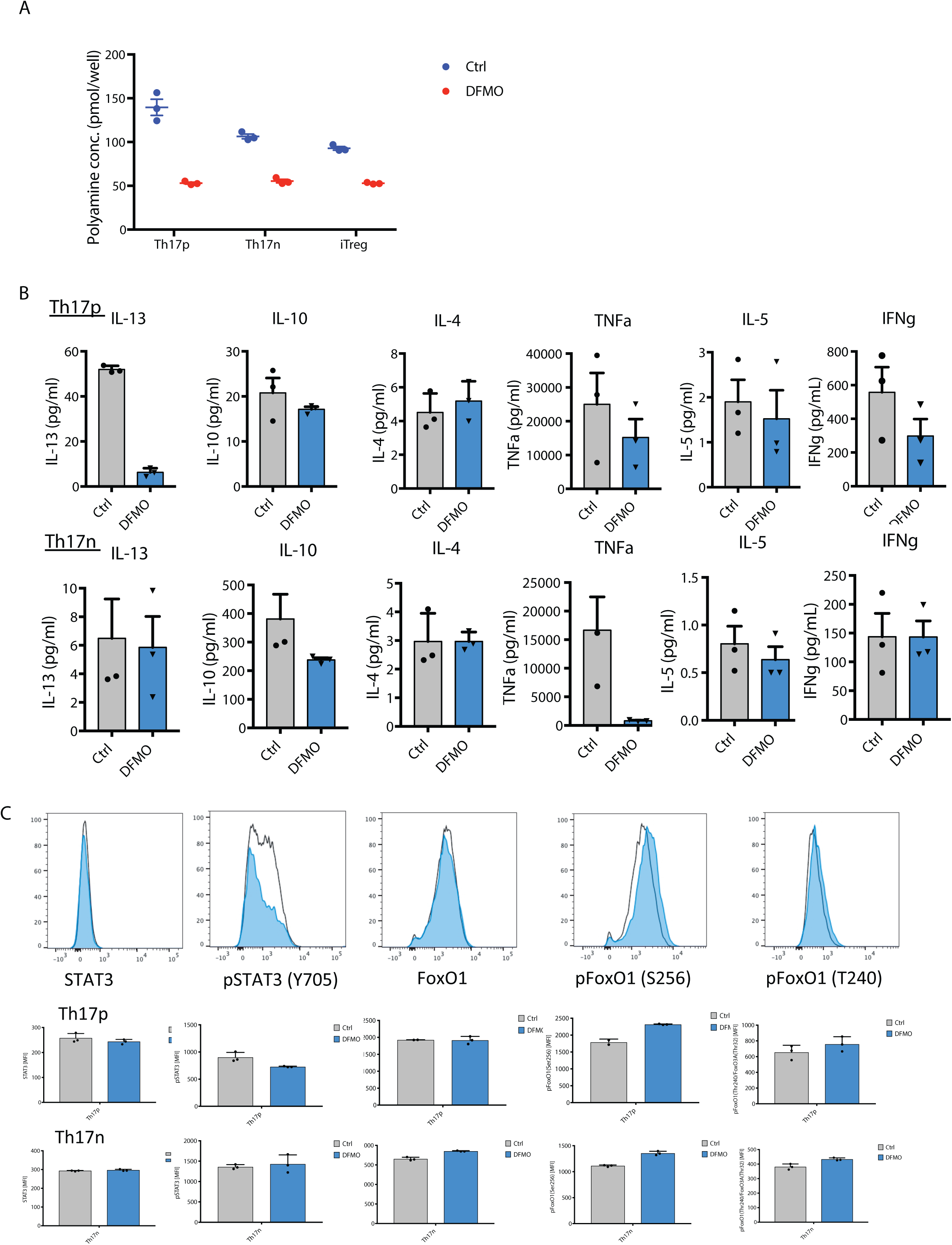
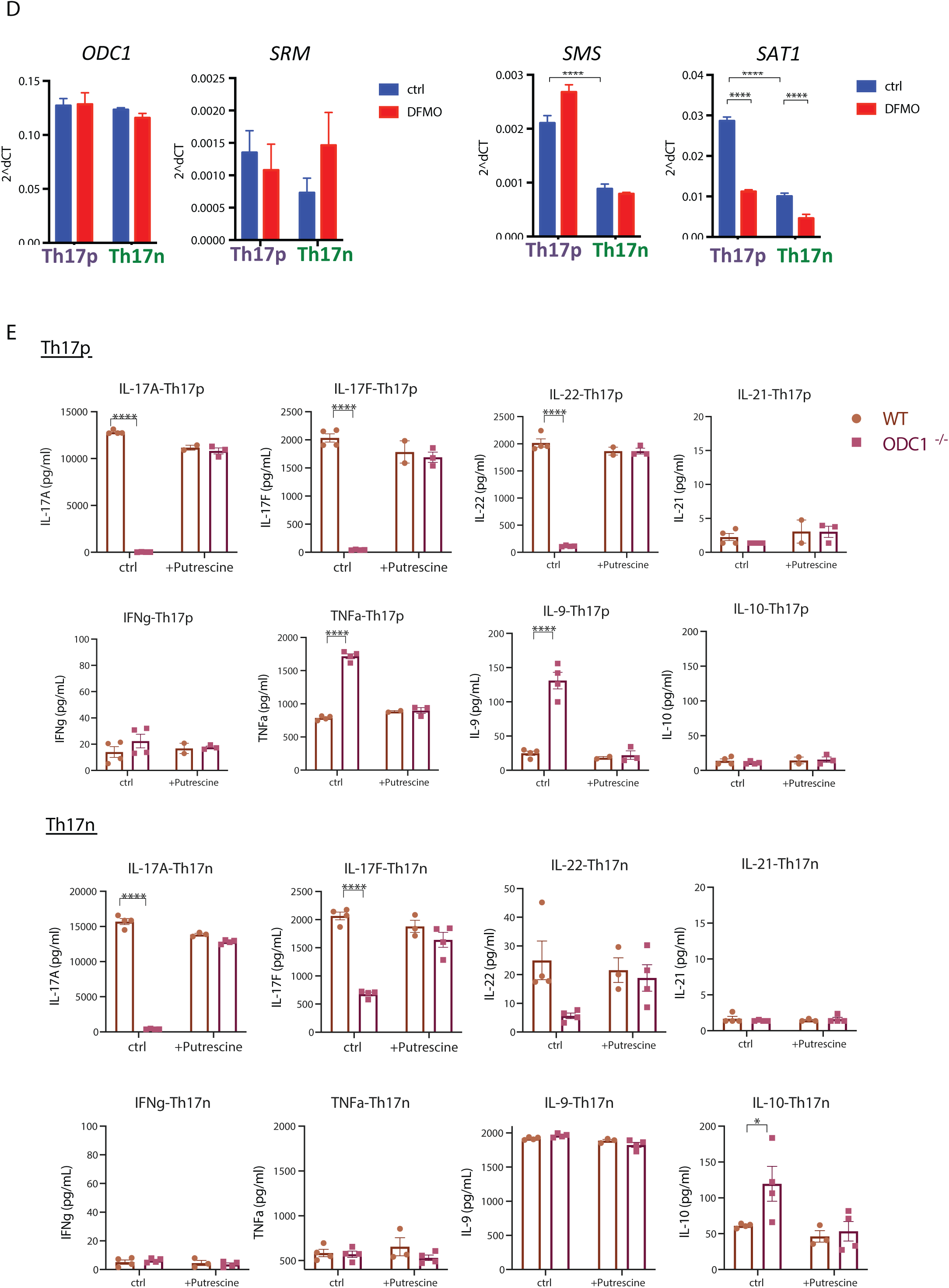
Chemical and genetic interference with the polyamine pathway suppress canonical Th17 cell cytokines. (**A**) The effect of DFMO on cellular polyamine concentration is measured by an enzymatic assay. Th17p, Th17n and iTregs are differentiated in the presence of DFMO and harvested at 96 hours for analysis. (**B**) Additional analysis of cytokines in supernatant as in Figure 2C. (**C**) Protein and phospho-protein analysis by flow cytometry for Th17n and Th17p cells treated with control of DFMO. (**D**) The effect of DFMO on enzymes in the polyamine pathway as measured by qPCR. Th17p and Th17n cells were differentiated in the presence of control or DFMO and harvested at 48h for RNA extraction and qPCR analysis. (**E**) The effect of genetic perturbation of ODC1 on cytokine production from Th17p (upper panels) and Th17n cells (lower panels). Supernatant from Th17p and Th17n differentiation culture was harvested at 96 hours and analyzed by legendplex for cytokine concentration.

**Supplemental Figure 3.**
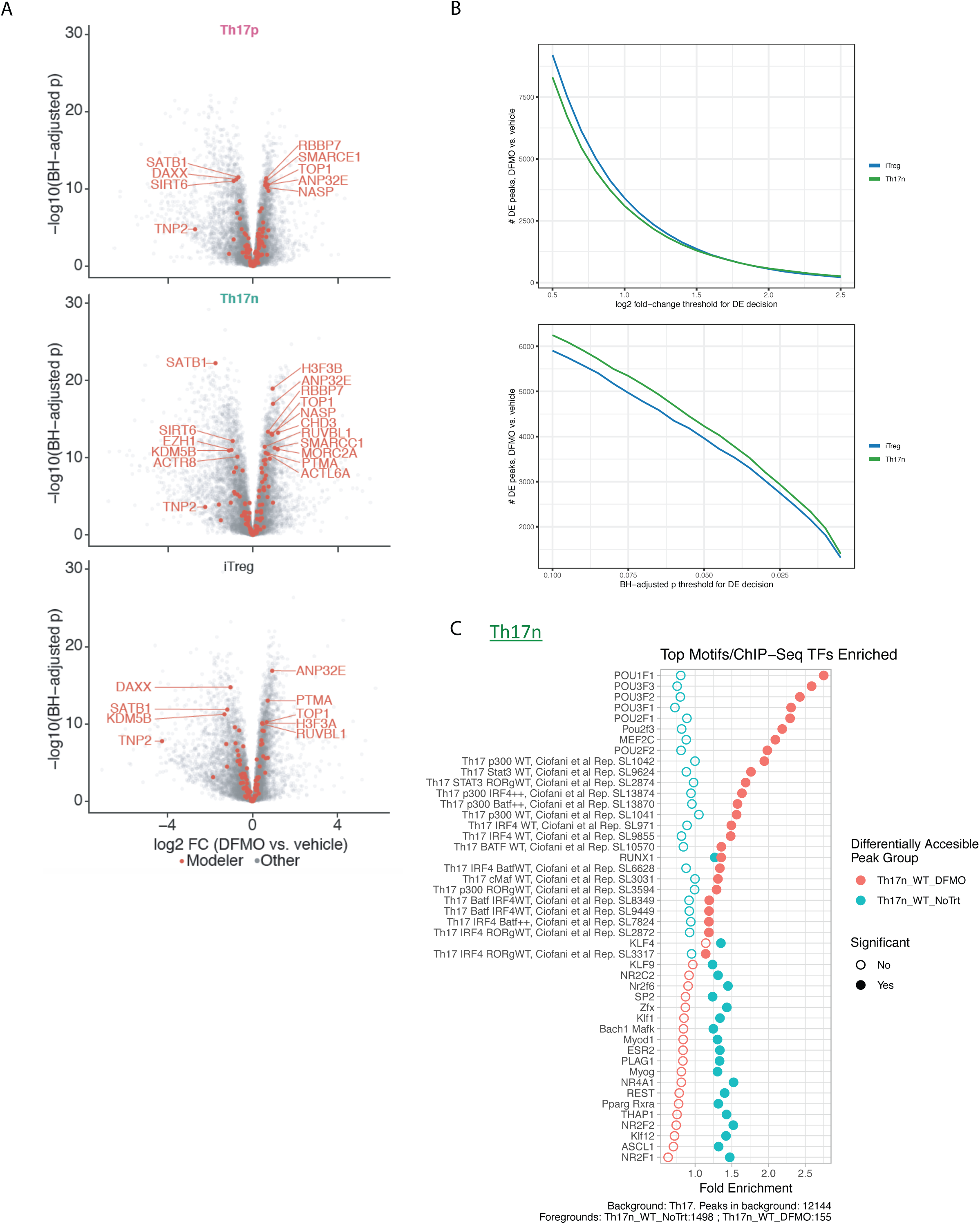
DFMO treatment promotes Treg-like transcriptome and epigenome. (**A**) Volcano plots showing affected chromatin modifiers by DFMO treatment in Th17n, Th17p and iTreg cells. (**B)** Number of differentially expressed (DE) peaks between DFMO and vehicle-treated cells as a function of the significance threshold. Upper panel, log2FC used as threshold; Lower panel, BH-adjusted P used as threshold. (**C**) Motif enrichment analysis of *in vitro* differentiated Th17n in the presence (red) or absence (green) of DFMO for Th17 specific genes.

**Supplemental Figure 4.**
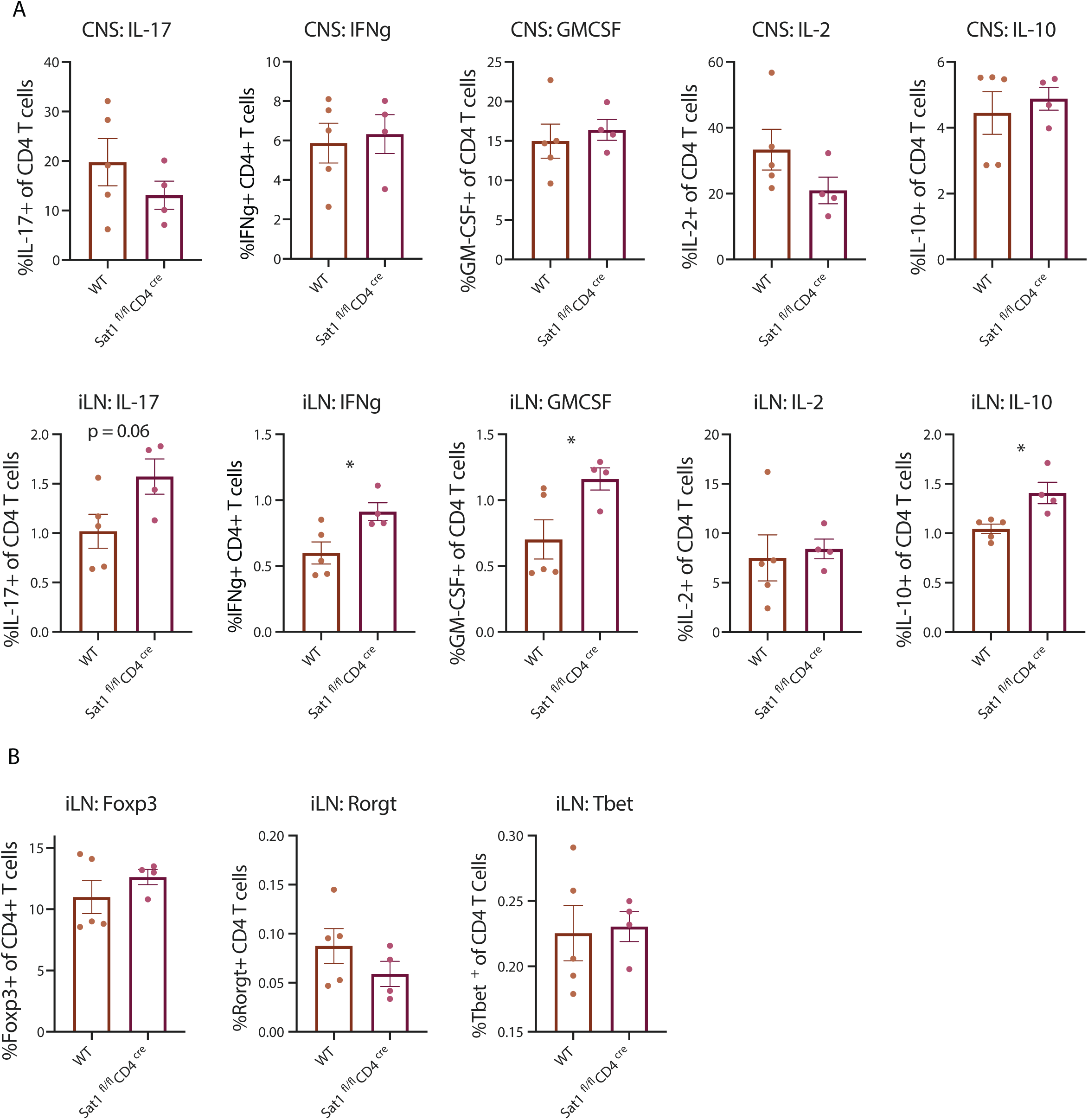
Targeting ODC1 and SAT1 alleviate EAE. Cells were isolated from CNS or inguinal lymph node of WT or SAT1^fl/fl^CD4^cre^ mice on day 15 post EAE induction (similar experiments as in Figure 4F). (**A**) Intracellular cytokines were measured by flow cytometry after 4-hour PMA/ionomycine stimulation ex vivo in the presence of brefaldin and monensin. (**B**) Transcription factors were analyzed directly ex vivo by intracellular staining.

## REFERENCE

[1] M. Kleinewietfeld and D. A. Hafler, “The plasticity of human Treg and Th17 cells and its role in autoimmunity,” Semin Immunol, vol. 25, pp. 305–12, Nov 15 2013.

[2] M. Noack and P. Miossec, “Th17 and regulatory T cell balance in autoimmune and inflammatory diseases,” Autoimmun Rev, vol. 13, pp. 668–77, Jun 2014.

[3] E. Bettelli, Y. Carrier, W. Gao, T. Korn, T. B. Strom, M. Oukka, et al., “Reciprocal developmental pathways for the generation of pathogenic effector TH17 and regulatory T cells,” Nature, vol. 441, pp. 235–8, May 11 2006.

[4] P. R. Mangan, L. E. Harrington, D. B. O’Quinn, W. S. Helms, D. C. Bullard, C. O. Elson, et al., “Transforming growth factor-beta induces development of the T(H)17 lineage,” Nature, vol. 441, pp. 231–4, May 11 2006.

[5] M. Veldhoen, R. J. Hocking, C. J. Atkins, R. M. Locksley, and B. Stockinger, “TGFbeta in the context of an inflammatory cytokine milieu supports de novo differentiation of IL-17-producing T cells,” Immunity, vol. 24, pp. 179–89, Feb 2006.

[6] M. J. McGeachy and D. J. Cua, “Th17 cell differentiation: the long and winding road,” Immunity, vol. 28, pp. 445–53, Apr 2008.

[7] L. Guglani and S. A. Khader, “Th17 cytokines in mucosal immunity and inflammation,” Curr Opin HIV AIDS, vol. 5, pp. 120–7, Mar 2010.

[8] W. Ouyang, J. K. Kolls, and Y. Zheng, “The biological functions of T helper 17 cell effector cytokines in inflammation,” Immunity, vol. 28, pp. 454–67, Apr 2008.

[9] E. Bettelli, T. Korn, M. Oukka, and V. K. Kuchroo, “Induction and effector functions of T(H)17 cells,” Nature, vol. 453, pp. 1051–7, Jun 19 2008.

[10] T. Korn, E. Bettelli, M. Oukka, and V. K. Kuchroo, “IL-17 and Th17 Cells,” Annu Rev Immunol, vol. 27, pp. 485–517, 2009.

[11] S. L. Gaffen, N. Hernandez-Santos, and A. C. Peterson, “IL-17 signaling in host defense against Candida albicans,” Immunol Res, vol. 50, pp. 181–7, Aug 2011.

[12] L. Romani, “Immunity to fungal infections,” Nat Rev Immunol, vol. 11, pp. 275–88, Apr 2011.

[13] A. Jager, V. Dardalhon, R. A. Sobel, E. Bettelli, and V. K. Kuchroo, “Th1, Th17, and Th9 effector cells induce experimental autoimmune encephalomyelitis with different pathological phenotypes,” J Immunol, vol. 183, pp. 7169–77, Dec 1 2009.

[14] Y. Lee, A. Awasthi, N. Yosef, F. J. Quintana, S. Xiao, A. Peters, et al., “Induction and molecular signature of pathogenic TH17 cells,” Nat Immunol, vol. 13, pp. 991–9, Oct 2012.

[15] M. J. McGeachy, K. S. Bak-Jensen, Y. Chen, C. M. Tato, W. Blumenschein, T. McClanahan, et al., “TGF-beta and IL-6 drive the production of IL-17 and IL-10 by T cells and restrain T(H)-17 cell-mediated pathology,” Nat Immunol, vol. 8, pp. 1390–7, Dec 2007.

[16] A. Awasthi, L. Riol-Blanco, A. Jager, T. Korn, C. Pot, G. Galileos, et al., “Cutting edge: IL-23 receptor gfp reporter mice reveal distinct populations of IL-17-producing cells,” J Immunol, vol. 182, pp. 5904–8, May 15 2009.

[17] D. J. Cua, J. Sherlock, Y. Chen, C. A. Murphy, B. Joyce, B. Seymour, et al., “Interleukin-23 rather than interleukin-12 is the critical cytokine for autoimmune inflammation of the brain,” Nature, vol. 421, pp. 744–8, Feb 13 2003.

[18] M. J. McGeachy, Y. Chen, C. M. Tato, A. Laurence, B. Joyce-Shaikh, W. M. Blumenschein, et al., “The interleukin 23 receptor is essential for the terminal differentiation of interleukin 17-producing effector T helper cells in vivo,” Nat Immunol, vol. 10, pp. 314–24, Mar 2009.

[19] Y. Chung, S. H. Chang, G. J. Martinez, X. O. Yang, R. Nurieva, H. S. Kang, et al., “Critical regulation of early Th17 cell differentiation by interleukin-1 signaling,” Immunity, vol. 30, pp. 576–87, Apr 17 2009.

[20] K. Ghoreschi, A. Laurence, X. P. Yang, C. M. Tato, M. J. McGeachy, J. E. Konkel, et al., “Generation of pathogenic T(H)17 cells in the absence of TGF-beta signalling,” Nature, vol. 467, pp. 967–71, Oct 21 2010.

[21] Y. Lee, M. Collins, and V. K. Kuchroo, “Unexpected targets and triggers of autoimmunity,” J Clin Immunol, vol. 34 Suppl 1, pp. S56–60, Jul 2014.

[22] C. E. Zielinski, F. Mele, D. Aschenbrenner, D. Jarrossay, F. Ronchi, M. Gattorno, et al., “Pathogen-induced human TH17 cells produce IFN-gamma or IL-10 and are regulated by IL-1beta,” Nature, vol. 484, pp. 514–8, Apr 26 2012.

[23] C. Wang, N. Yosef, J. Gaublomme, C. Wu, Y. Lee, C. B. Clish, et al., “CD5L/AIM Regulates Lipid Biosynthesis and Restrains Th17 Cell Pathogenicity,” Cell, vol. 163, pp. 1413–27, Dec 3 2015.

[24] J. T. Gaublomme, N. Yosef, Y. Lee, R. S. Gertner, L. V. Yang, C. Wu, et al., “Single-Cell Genomics Unveils Critical Regulators of Th17 Cell Pathogenicity,” Cell, vol. 163, pp. 1400–12, Dec 3 2015.

[25] J. D. Orth, I. Thiele, and B. O. Palsson, “What is flux balance analysis?,” Nat Biotechnol, vol. 28, pp. 245–8, Mar 2010.

[26] E. J. O’Brien, J. M. Monk, and B. O. Palsson, “Using Genome-scale Models to Predict Biological Capabilities,” Cell, vol. 161, pp. 971–987, May 21 2015.

[27] I. Thiele, N. Swainston, R. M. Fleming, A. Hoppe, S. Sahoo, M. K. Aurich, et al., “A community-driven global reconstruction of human metabolism,” Nat Biotechnol, vol. 31, pp. 419–25, May 2013.

[28] T. L. Bowlin, B. J. McKown, and P. S. Sunkara, “The effect of alpha-difluoromethylornithine, an inhibitor of polyamine biosynthesis, on mitogen-induced interleukin 2 production,” Immunopharmacology, vol. 13, pp. 143–7, Apr 1987.

[29] J. Jell, S. Merali, M. L. Hensen, R. Mazurchuk, J. A. Spernyak, P. Diegelman, et al., “Genetically altered expression of spermidine/spermine N1-acetyltransferase affects fat metabolism in mice via acetyl-CoA,” J Biol Chem, vol. 282, pp. 8404–13, Mar 16 2007.

[30] A. E. Pegg, “Spermidine/spermine-N(1)-acetyltransferase: a key metabolic regulator,” Am J Physiol Endocrinol Metab, vol. 294, pp. E995–1010, Jun 2008.

[31] B. C. Mounce, E. Z. Poirier, G. Passoni, E. Simon-Loriere, T. Cesaro, M. Prot, et al., “Interferon-Induced Spermidine-Spermine Acetyltransferase and Polyamine Depletion Restrict Zika and Chikungunya Viruses,” Cell Host Microbe, vol. 20, pp. 167–77, Aug 10 2016.

[32] S. Xiao, N. Yosef, J. Yang, Y. Wang, L. Zhou, C. Zhu, et al., “Small-molecule RORgammat antagonists inhibit T helper 17 cell transcriptional network by divergent mechanisms,” Immunity, vol. 40, pp. 477–89, Apr 17 2014.

[33] Q. Li, J. Zou, M. Wang, X. Ding, I. Chepelev, X. Zhou, et al., “Critical role of histone demethylase Jmjd3 in the regulation of CD4+ T-cell differentiation,” Nat Commun, vol. 5, p. 5780, 2014.

[34] E. Karouzakis, R. E. Gay, S. Gay, and M. Neidhart, “Increased recycling of polyamines is associated with global DNA hypomethylation in rheumatoid arthritis synovial fibroblasts,” Arthritis Rheum, vol. 64, pp. 1809–17, Jun 2012.

[35] H. C. Hsu, J. R. Seibold, and T. J. Thomas, “Regulation of ornithine decarboxylase in the kidney of autoimmune mice with the lpr gene,” Autoimmunity, vol. 19, pp. 253–64, 1994.

[36] W. H. Brooks, “Increased polyamines alter chromatin and stabilize autoantigens in autoimmune diseases,” Front Immunol, vol. 4, p. 91, 2013.

[37] A. E. Pegg, “Mammalian polyamine metabolism and function,” IUBMB Life, vol. 61, pp. 880–94, Sep 2009.

[38] A. E. Pegg, “Functions of Polyamines in Mammals,” J Biol Chem, vol. 291, pp. 14904–12, Jul 15 2016.

[39] D. Kraus, Q. Yang, D. Kong, A. S. Banks, L. Zhang, J. T. Rodgers, et al., “Nicotinamide N-methyltransferase knockdown protects against diet-induced obesity,” Nature, vol. 508, pp. 258–62, Apr 10 2014.

[40] A. C. Childs, D. J. Mehta, and E. W. Gerner, “Polyamine-dependent gene expression,” Cell Mol Life Sci, vol. 60, pp. 1394–406, Jul 2003.

[41] K. Tamari, M. Konno, A. Asai, J. Koseki, K. Hayashi, K. Kawamoto, et al., “Polyamine flux suppresses histone lysine demethylases and enhances ID1 expression in cancer stem cells,” Cell Death Discov, vol. 4, p. 104, 2018.

[42] T. Murray-Stewart, M. Dunworth, J. R. Foley, C. E. Schwartz, and R. A. Casero, Jr., “Polyamine Homeostasis in Snyder-Robinson Syndrome,” Med Sci (Basel), vol. 6, Dec 7 2018.

[43] D. Cipolletta, D. Kolodin, C. Benoist, and D. Mathis, “Tissular T(regs): a unique population of adipose-tissue-resident Foxp3+CD4+ T cells that impacts organismal metabolism,” Semin Immunol, vol. 23, pp. 431–7, Dec 2011.

[44] S. Epelman, K. J. Lavine, and G. J. Randolph, “Origin and functions of tissue macrophages,” Immunity, vol. 41, pp. 21–35, Jul 17 2014.

[45] N. E. Lewis, H. Nagarajan, and B. O. Palsson, “Constraining the metabolic genotype-phenotype relationship using a phylogeny of in silico methods,” Nat Rev Microbiol, vol. 10, pp. 291–305, Feb 27 2012.

[46] B. O. Palsson, Systems Biology: Constraint-Based Reconstruction and Analysis. 2nd ed.edition. : Cambridge University Press., 2015.

[47] M. Singer, C. Wang, L. Cong, N. D. Marjanovic, M. S. Kowalczyk, H. Zhang, et al., “A Distinct Gene Module for Dysfunction Uncoupled from Activation in Tumor-Infiltrating T Cells,” Cell, vol. 166, pp. 1500–1511 e9, Sep 08 2016.

[48] M. B. Cole, D. Risso, A. Wagner, D. DeTomaso, J. Ngai, E. Purdom, et al., “Performance Assessment and Selection of Normalization Procedures for Single-Cell RNA-Seq,” Cell Syst, vol. 8, pp. 315–328 e8, Apr 24 2019.

[49] M. R. Corces, J. D. Buenrostro, B. Wu, P. G. Greenside, S. M. Chan, J. L. Koenig, et al., “Lineage-specific and single-cell chromatin accessibility charts human hematopoiesis and leukemia evolution,” Nat Genet, vol. 48, pp. 1193–203, Oct 2016.

[50] A. M. Bolger, M. Lohse, and B. Usadel, “Trimmomatic: a flexible trimmer for Illumina sequence data,” Bioinformatics, vol. 30, pp. 2114–20, Aug 1 2014.

[51] B. Langmead and S. L. Salzberg, “Fast gapped-read alignment with Bowtie 2,” Nat Methods, vol. 9, pp. 357–9, Mar 4 2012.

[52] J. D. Buenrostro, P. G. Giresi, L. C. Zaba, H. Y. Chang, and W. J. Greenleaf, “Transposition of native chromatin for fast and sensitive epigenomic profiling of open chromatin, DNA-binding proteins and nucleosome position,” Nat Methods, vol. 10, pp. 1213–8, Dec 2013.

[53] Y. Zhang, T. Liu, C. A. Meyer, J. Eeckhoute, D. S. Johnson, B. E. Bernstein, et al., “Model-based analysis of ChIP-Seq (MACS),” Genome Biol, vol. 9, p. R137, 2008.

[54] M. I. Love, W. Huber, and S. Anders, “Moderated estimation of fold change and dispersion for RNA-seq data with DESeq2,” Genome Biol, vol. 15, p. 550, 2014.

[55] A. Khan, O. Fornes, A. Stigliani, M. Gheorghe, J. A. Castro-Mondragon, R. van der Lee, et al., “JASPAR 2018: update of the open-access database of transcription factor binding profiles and its web framework,” Nucleic Acids Res, vol. 46, pp. D260–D266, Jan 4 2018.

[56] C. E. Grant, T. L. Bailey, and W. S. Noble, “FIMO: scanning for occurrences of a given motif,” Bioinformatics, vol. 27, pp. 1017–8, Apr 1 2011.

[57] R. Lesurf, K. C. Cotto, G. Wang, M. Griffith, K. Kasaian, S. J. Jones, et al., “ORegAnno 3.0: a community-driven resource for curated regulatory annotation,” Nucleic Acids Res, vol. 44, pp. D126–32, Jan 4 2016.

[58] C. Y. McLean, D. Bristor, M. Hiller, S. L. Clarke, B. T. Schaar, C. B. Lowe, et al., “GREAT improves functional interpretation of cis-regulatory regions,” Nat Biotechnol, vol. 28, pp. 495–501, May 2010.

